# The Vomeronasal System of *Talpa occidentalis*: A Combined Histological, Immunohistochemical, and Lectin-Binding Approach

**DOI:** 10.1101/2025.06.26.661819

**Authors:** Gadea García Hernando, Antía Martínez Antonio, Mostafa G. A. Elsayed, Alex Vázquez Castiñeira, Pablo Sanchez Quinteiro, Irene Ortiz Leal

## Abstract

The vomeronasal system (VNS) is critical for detecting pheromonal cues that modulate sociosexual behaviors. Despite its central role in chemical communication, our understanding of its anatomical and functional variability across mammals remains incomplete. This study provides the first detailed characterization of the VNS in the Iberian mole (*Talpa occidentalis*), a fossorial species endemic to the Iberian Peninsula. We performed a morphofunctional and neurochemical analysis of the vomeronasal organ (VNO) and the accessory olfactory bulb (AOB) using histology, immunohistochemistry, and lectin histochemistry. The VNO in *T. occidentalis* exhibited an unusual circular lumen lined by a uniform sensory epithelium, lacking the dual epithelial organization seen in most species. The vomeronasal cartilage was limited in extent and did not form the typical J-shaped structure. Importantly, no evidence of a vomeronasal pump was found, suggesting alternative mechanisms for semiochemical entry, likely facilitated by the organ’s anatomical position and continuous receptor distribution. Immunohistochemical analysis revealed strong expression of Gαi2 and Gϒ8 in sensory neurons, with weaker Gα0 expression, suggesting predominance of V1R-type signal transduction. The AOB, though small, exhibited clear lamination and specific marker localization (Gαi2, OMP, CR, MAP2), indicating robust functional organization. Lectin binding revealed specific glycosylation patterns in the glomerular layer, with STL and LEA marking synaptic regions. These findings uncover unprecedented anatomical and molecular features in the VNS of *T. occidentalis*, positioning this species as a valuable model for studying vomeronasal diversity and evolution among Laurasiatherian mammals.

## Introduction

Chemical communication is a fundamental mechanism across animal taxa, mediating key processes such as reproduction, social interaction, foraging, and predator avoidance (Wyatt, 2013). Semiochemicals—volatile or non-volatile molecules that carry information between individuals—act either within species (pheromones) or across species boundaries (allelochemicals), and their detection is crucial for species survival. In terrestrial environments, semiochemicals tend to be small, volatile compounds, whereas aquatic species rely on highly soluble molecules, often conjugated with sulfate or glucuronic acid to enhance signal dispersal (Mollo et al., 2017; Bowers et al., 2023). Pheromones, in particular, play a central role in coordinating reproductive behavior, maternal bonding, territoriality, and kin recognition (Hayes et al., 2003; Baum and Cherry, 2015; Mota-Rojas et al., 2024). Their detection is mediated by two complementary chemosensory systems: the main olfactory system and the vomeronasal system (VNS), each with distinct anatomical, molecular, and functional characteristics (Halpern and Martínez-Marcos, 2003; Tirindelli et al., 2009; Salazar et al., 2016; Kobayashi-Sakashita et al., 2023).

The vomeronasal system (VNS) comprises three main components: the vomeronasal organ (VNO), the vomeronasal nerves, and the accessory olfactory bulb (AOB). The VNO is a paired, elongated tubular structure located bilaterally at the base of the nasal septum, enclosed in cartilage or bone depending on the species (Ignacio Salazar et al., 1998). It is lined by a pseudostratified neuroepithelium composed of three distinct cell layers: a superficial layer of sustentacular cells, an intermediate layer containing bipolar vomeronasal receptor neurons (VRNs), and a basal layer of progenitor cells (Adams, 1972). These VRNs extend apical dendrites toward the lumen to detect pheromonal cues and project their axons posteriorly to form the vomeronasal nerves. These nerves travel dorsocaudally beneath the nasal mucosa, pass through the cribriform plate of the ethmoid bone, and terminate in the glomerular layer of the AOB (Mori et al., 1987). The AOB, typically situated on the dorsomedial aspect of the olfactory peduncle, is the first central processing station for vomeronasal input (Meisami and Bhatnagar, 1998; Ortiz-Leal et al., 2022b). It shares a laminar organization with the main olfactory bulb (MOB), comprising a nerve layer, a glomerular layer, a mitral-plexiform layer, and a deep granular layer (Ortiz-Leal et al., 2022a). However, notable differences include the smaller size and less defined boundaries of vomeronasal glomeruli, and the fact that mitral cells in the AOB often extend dendrites to multiple glomeruli, unlike the one-to-one mapping seen in the MOB (Larriva-Sahd, 2008; Ruiz-Rubio et al., 2024a). Information processed in the AOB is relayed to limbic and hypothalamic structures— such as the medial amygdala, the bed nucleus of the stria terminalis, and various hypothalamic nuclei—without engaging higher-order cortical centers. This architecture supports the VNS’s role in eliciting instinctive, rapid behavioral and neuroendocrine responses to socially relevant chemical signals (Wu et al., 2014; Nguyen et al., 2024).

Due to its species-specific anatomical adaptations, the VNO exhibits remarkable interspecific variability, precluding the extrapolation of data across taxa. Comparative studies have shown that in some mammalian clades the VNO is well developed, while in others it is regressed or entirely absent (Villamayor et al., 2021; Torres et al., 2023b). Thus, each species must be studied independently, especially those with unique ecological and sensory constraints (Salazar and Sanchez Quinteiro, 2009). From a phylogenetic and molecular perspective, much of this variation arises from differences in the expression of two families of vomeronasal receptors: V1Rs and V2Rs, both G protein-coupled receptors (GPCRs), yet distinct in ligand affinity, signaling mechanisms, and anatomical organization (Murata et al., 2024a). V1Rs are typically found in apically located VRNs, which project to the anterior AOB and couple to Gαi2 proteins, detecting mainly small, volatile compounds (Torres et al., 2022; Weiss et al., 2023). Conversely, V2Rs are expressed by basal VRNs, project to the posterior AOB, and signal via Gαo proteins; these receptors detect larger, peptide- or protein-based ligands, including major urinary proteins and pathogen-derived (Silvotti et al., 2007; Ortiz-Leal et al., 2024).

The proportion and functionality of these receptor types vary considerably across species. For instance, murine models exhibit robust expression of both receptor families, reflecting their heavy reliance on chemical communication (Jia and Halpern, 1996). In contrast, several carnivores and primates exhibit reduced or absent V2R repertoires (Rodriguez and Mombaerts, 2002; Ortiz-Leal et al., 2024), and in some cases, the VNO itself is vestigial (Kondoh et al., 2024). These evolutionary trends suggest that ecological pressures—such as nocturnality, fossoriality, or solitary behavior—selectively shape vomeronasal receptor expression and the degree of functional specialization. In this context, studying the vomeronasal system of fossorial mammals such as the Iberian mole (*Talpa occidentalis*) becomes especially relevant. Given their limited visual and auditory capabilities, these animals likely depend heavily on chemosensory input to navigate their environment, locate prey, avoid predators, and identify conspecifics (Macdonald et al., 1997). Nevertheless, to our knowledge no data are currently available for any member of the *Talpidae* family, making this a critical knowledge gap in the comparative anatomy and neurobiology of mammalian chemosensory systems.

The Iberian mole (*Talpa occidentalis*), a fossorial eulipotyphlan endemic to the Iberian Peninsula, represents a phylogenetically and ecologically distinct model for exploring vomeronasal function. Within the order Eulipotyphla —which includes hedgehogs (Erinaceidae), shrews (Soricidae), and moles (Talpidae) (Douady et al., 2002)—only a few studies have characterized the VNS, notably in the hedgehog (*Atelerix albiventris*) (Kondoh et al., 2021) and the musk shrew (*Suncus murinus*) (Oikawa et al., 1993), highlighting the lack of data for *Talpa* spp. *Eulipotyphla* constitutes the earliest-diverging lineage within *Laurasiatheria*. This basal placement suggests that moles, and their close relatives, retain key ancestral traits and may offer valuable insights into the early evolution of boreoeutherian mammals (Zachos, 2020). Their phylogenetic proximity to the root of *Laurasiatheria* also makes them particularly relevant for comparative studies with *Euarchontoglires*, the sister superorder comprising primates, rodents, and lagomorphs (Meredith et al., 2011). Notably, both rodents and lagomorphs retain functional repertoires of vomeronasal receptors from both the V1R and V2R gene families, a trait that has been partially or completely lost in other mammalian lineages, such as primates and some carnivores (Suárez et al., 2011a; Ortiz-Leal et al., 2023). Understanding how the expression of these receptor families evolved—and in particular, how and when the V2R family was reduced or lost in certain clades—requires phylogenetically informed comparisons with basal laurasiatherian taxa. In this context, insectivorous species such as moles may offer key insights into the ancestral state of the vomeronasal system in *Boreoeutheria*.

To date, no histological or neuroanatomical studies of the VNS have been published for any *Talpidae* species. The current study addresses this gap by characterizing the vomeronasal system of *T. occidentalis* using an integrative morphofunctional approach, combining classical histology, immunohistochemistry, and lectin histochemistry. Through this multimodal analysis, we aim to: (i) describe the microanatomical features of the VNO and AOB; (ii) identify specific glycosylation patterns and molecular markers associated with vomeronasal sensory neurons; and (iii) infer the potential functional organization of the system in the context of a fossorial mammal. Our findings will not only expand the anatomical and neurochemical knowledge of the vomeronasal system in Eulipotyphla, but also provide a comparative framework for future studies on chemosensory specialization in mammals with extreme ecological niches.

## Methods

### Origin of the samples

A total of five Iberian moles (*Talpa occidentalis*) were used in this study. All specimens belonged to the sample collection of the Anatomy Unit at the Faculty of Veterinary Medicine, Campus Terra, University of Santiago de Compostela (Lugo, Spain). All specimens were preserved in Bouin’s solution, except for one specimen, which was fixed in 10% neutral buffered formalin.

### Sample extraction

The study focused on the following anatomical structures: nasal cavity, vomeronasal organ, accessory and main olfactory bulbs.

#### Nasal cavity

The objective of nasal cavity processing was to perform a comprehensive microscopic analysis of the vomeronasal organ at different rostrocaudal levels. In addition, the study aimed to identify the incisive and vomeronasal ducts, as well as the communication between them. To enable microscopic examination, decalcification was necessary to soften the bony structures and prevent tissue rupture during microtome sectioning. Specimens were immersed in a decalcifying solution (Osteomoll, Sigma) under continuous agitation at room temperature for 2 days. Following standard dehydration and clearing procedures, the nasal cavities were embedded in paraffin, resulting in five blocks that were serially sectioned in the transverse plane at 7 µm thickness using a rotary microtome.

#### Olfactory bulbs (OB)

Whole-brain specimens were processed as single paraffin blocks to enable general morphological assessment. To extract the brain from the skull, the cranial bones were carefully removed using a bone rongeur. The rostral part of the telencephalon, containing the olfactory bulbs, was dissected and paraffin embedded. Sections were obtained using a rotary microtome in both sagittal and transverse planes, in order to maximize anatomical information.

### Paraffin Embedding

To prepare the specimens for paraffin embedding, they were first subjected to a dehydration process to remove water from the tissues, enabling proper infiltration of liquid paraffin into cellular and interstitial spaces. Dehydration began with immersion in 70% ethanol for 2 hours, followed by 90% ethanol for 1 hour, and then 96% ethanol for 2 hours. This was followed by three consecutive baths in 100% ethanol, each lasting 1 hour. The tissues were then placed in an ethanol–xylene solution for 1 hour and 15 minutes, followed by two clearing steps in pure xylene: the first for 1 hour and 30 minutes, and the second for 45 minutes.

### General histological stains

For the realization of this study, we have used the following stains:

#### Hematoxylin-eosin (H-E)

This staining method was chosen to allow general histological assessment of the different structures, as it offers good contrast and clear visualization of cellular and tissue architecture. Hematoxylin, a basic dye, selectively stains nuclei, whereas eosin, an acidic dye, highlights the cytoplasm and extracellular components.

#### Nissl Staining

Cresyl violet staining is particularly suited for the examination of nervous tissue, as it binds selectively to nucleic acid-rich structures, including the nucleus, free cytoplasmic ribosomes, and rough endoplasmic reticulum. This property allows for detailed visualization of neuronal somata and glial cells, as well as the proximal segments of neuronal processes. The technique is widely employed to evaluate the organization, distribution, and density of cells within neural tissues.

#### PAS

The periodic acid–Schiff (PAS) technique is particularly useful for highlighting glycoconjugates in tissue sections, especially those rich in carbohydrates such as glycoproteins, glycolipids, and mucopolysaccharides. By reacting with periodic acid, these components are oxidized to aldehydes, which subsequently bind to the Schiff reagent, producing a distinctive magenta coloration. This staining method is frequently employed to visualize basement membranes, neutral mucins, and other carbohydrate-rich structures, offering valuable insight into the distribution and organization of extracellular matrix elements. The protocol followed is detailed in depth in Salazar et al. (1998).

#### Alcian blue (AB)

This basic dye is used to stain acidic substances, primarily acidic mucopolysaccharides. It binds to carbohydrates and is used in combination with PAS staining to differentiate between acidic and neutral mucins. Alcian Blue is a cationic dye that binds to anionic groups of acidic mucins, producing a turquoise-blue color. This staining is particularly effective in glandular tissues, as well as in structural components such as cartilage and bone. The detailed protocol applied is specified in Torres et al. (2021).

#### Gallego’s Trichrome Staining

This staining technique is used to differentiate the components of connective tissue and muscle. It employs three dyes—two cytoplasmic and one nuclear—that stain collagen blue, cartilage fuchsia, nuclei purple, and muscle fibers green, providing strong contrast between tissue structures. The protocol followed is detailed in depth in Torres et al. (2023a).

### Immunohistochemical staining

To examine the distribution of specific neuronal markers and G protein subunits, immunohistochemical staining was conducted using an indirect method based on peroxidase labeling. This approach takes advantage of the selective interaction between antigens and antibodies, allowing precise localization of target proteins in paraffin-embedded tissue sections under the microscope. Prior to antibody incubation, tissue sections were deparaffinized in xylene and passed through a descending ethanol gradient into 0.1 M phosphate buffer (PB), a rehydration step necessary to restore near-physiological osmotic conditions. Endogenous peroxidase activity, which could otherwise lead to nonspecific staining, was quenched by incubating the sections in a 3% hydrogen peroxide (H₂O₂) solution prepared in distilled water. To further minimize background and enhance specificity, sections were pretreated with 2% bovine serum albumin (BSA) in 0.1 M PB to block nonspecific protein-binding sites. The staining protocol followed a two-step immunohistochemical procedure:

Tissue sections were incubated overnight at 4 °C with the corresponding primary antibody to allow specific binding to the target antigen. The following day, sections were exposed for 30 minutes at room temperature to a horseradish peroxidase (HRP)-conjugated secondary antibody polymer (CRF Anti-Polyvalent HRP Polymer; ScyTek, Cache Valley, USA), designed to bind to the primary antibody and amplify the signal. After both incubation steps, any unbound antibodies were thoroughly removed through multiple washes in phosphate buffer (PB), followed by a final rinse in 0.1 M TRIS buffer to ensure optimal conditions for chromogen development. Immunolabeling was visualized by applying 3,3′-diaminobenzidine (DAB) in the presence of hydrogen peroxide. The enzymatic activity of peroxidase catalyzed DAB oxidation, resulting in a localized brown precipitate at immunoreactive sites. Sections were subsequently rinsed in PB and distilled water, and then coverslipped using a permanent mounting medium suitable for brightfield microscopy. Positive controls consisted of tissue samples known to express the antigen of interest, while negative controls were processed identically but without the primary antibody, ensuring the specificity of the staining procedure.

**Table 1.** Primary antibodies. The following primary antibodies were used

| Antibody | Type | Species | Inmunogen | Dilution | Catalog No. |
| --- | --- | --- | --- | --- | --- |
| Anti-Gai2 | PC | Rabbit | Peptide mapping within a highly divergent domain of rat Gai2 | 1:150 | ProteinTech 11136-1-AP |
| Anti-Gao | PC | Rabbit | Gao subunit of the bovine GTP-binding protein | 1:100 | Santa Cruz SC-387 |
| Anti-GY8 | PC | Rabbit | Recombinant GNg8 expresado en *E. coli* | 1:150 | Cloud-Clone. PAQ769Mu01 |
| Anti-GAP43 | PC | Rabbit | - | 1:200 | ProteinTech 16971-1-AP |
| Anti-OMP | MC | Mouse | Human OMP Aminoacid 1-163 | 1:200 | Santa Cruz SC-365818 |
| Anti-GFAP |  | Rabbit | - | 1:300 | Dako Z0334 |
| Anti-MAP-2 |  | Mouse | - | 1:200 | Sigma G9264 |
| Anti-CB | PC | Rabbit | Recombinant rat calbindin D-28k | 1:200 | ProteinTech 14479-1-AP |
| Anti-CR | PC | Rabbit | Recombinant human calretinin containing an N-terminal 6×His tag | 1:200 | ProteinTech 12278-1-AP |
| Anti-PGP | PC | Rabbit | UCHL1/PGP9.5 fusion protein Ag6490 | 1:200 | ProteinTech 14730-1-AP |
| Anti-Enolase | MC | Mouse | Amino acids 416–433 of human γ-enolase | 1:100 | Santa Cruz SC-21738 |
| Anti-NRP1 | PC | Rabbit | Recombinant NRP1 expressed in *E. coli* | 1:100 | Cloude-Clone Corp PAA692Hu01 |
| Anti-GNRH1 | PC | Rabbit | Human GNRH fusion protein | 1:200 | Elabscience E-AB-52158 |
| Anti-TUBb3 | PC | Rabbit | Recombinant TUBb3 expressed in *E. coli* | 1:100 | Cloude-Clone Corp PAE711Hu01 |
| Anti-NF-H | PC | Rabbit | Human NFH derived peptide | 1:200 | ELK Biotechnology ES6357 |
| Anti- Ki-67 | PC | Rabbit | Recombinant Ki67 expressed in *E. coli* | 1:100 | Cloude-Clone Corp<br>PACo47Hu01 |
| Anti-NSE | PC | Rabbit | Recombinant NSE expressed in *E. coli* | 1:100 | Cloude-Clone Corp<br>PAA537Mu01 |
| Anti-Robo 2 | PC | Rabbit | Synthetic peptide derived from human Robo2 | 1:200 | BT LAB BT-APo7965 |
| Anti-s100 | PC | Rabbit | Recombinant S100 expressed in *E. coli* | 1:100 | Cloude-Clone Corp<br>PAA012Hu01 |
| Anti-CK 20 | PC | Rabbit | Recombinant CK20 expressed in *E. coli* | 1:100 | Cloude-Clone Corp<br>PAB24oHu01 |
| Anti-CALB | PC | Rabbit | Recombinant CALB expressed in *E. coli* | 1:100 | Cloude-Clone Corp<br>PAG438Hu01 |
| Anti- SP | PC | Rabbit | Synthetic peptide from SP conjugated to OVA | 1:100 | Cloude-Clone Corp<br>PAA393Ra01 |
| Anti-NPY | PC | Rabbit | Recombinant NPY expressed in *E. coli* | 1:100 | Cloude-Clone Corp<br>PAA879Ra01 |

### Lectin histochemical labelling

Lectins are a class of glycan-recognizing proteins, also known as agglutinins, that bind selectively and reversibly to specific carbohydrate moieties through non-covalent interactions, without inducing chemical alterations in the glycoconjugates. They are typically isolated from a wide range of biological sources—most notably plants—and are commonly named after their organism of origin. (Plendl and Sinowatz, 1998). In this study, a panel of ten lectins was used to investigate the glycoconjugate composition of various olfactory and vomeronasal structures. Each lectin was applied to all tissue samples at working dilutions previously optimized through pilot experiments. The lectins employed, along with their sources, carbohydrate-binding preferences, and established specificities, are detailed below.

<u>STL (*Solanum tuberosum* lectin):</u> binds N-acetylglucosamine (GlcNAc) and its oligomers (McCurrach and Kilpatrick, 1986) .

<u>LEA (*Lycopersicon esculentum* agglutinin):</u> recognizes GlcNAc-rich N-glycans (Tomiyasu et al., 2018).

<u>DBA (*Dolichos biflorus* agglutinin):</u> shows high affinity for α-GalNAc, particularly terminal residues of blood group A antigens (Chun et al., 2024).

<u>SBA (*Glycine max* agglutinin):</u> binds primarily to β-galactose (Taniguchi et al., 1993) .

<u>VVA (*Vicia villosa* agglutinin):</u> targets D-galactose and N-acetylgalactosamine (GalNAc) (Shapiro et al., 1995).

<u>UEA (*Ulex europaeus* agglutinin):</u> binds terminal L-fucose; selectively labels olfactory and vomeronasal components (Kondoh et al., 2017a).

<u>ECL (*Erythrina cristagalli* lectin):</u> binds D-galactose and GlcNAc residues (Keller et al., 2022).

<u>WGA (*Triticum vulgaris* lectin):</u> binds GlcNAc and sialic acid (Lee et al., 2012).

<u>PHL (*Phaseolus lunatus* lectin)</u>: recognizes mannose residues, and to a lesser extent, glucose (Shin et al., 2017).

<u>LCA (*Lens culinaris* agglutinin)</u>: binds D-galactose residues (Salazar et al., 2008).

### Labeling procedure

Slides were first deparaffinized and rehydrated, followed by inhibition of endogenous tissue peroxidase activity by incubating the sections in 3% hydrogen peroxide (H₂O₂) for 10 minutes to prevent interference during the detection step. Nonspecific binding sites were then blocked with 2% bovine serum albumin (BSA) for 30 minutes. Subsequently, the sections were incubated overnight at 4 °C in a humidified chamber with biotinylated lectin diluted to 0.05%. On the following day, samples were incubated for 90 minutes at room temperature with the avidin–biotin–peroxidase complex (ABC), which binds to the lectin and enhances the detection reaction. Finally, immunoreactive sites were visualized by incubating the sections in a chromogenic solution containing 0.05% 3,3′-diaminobenzidine (DAB) and 0.003% H₂O₂ in 0.2 M Tris-HCl buffer, resulting in a brown precipitate at binding sites. The enzymatic reaction was monitored under a light microscope.

### Immunohistochemistry for Optical Microscopy in Free-floating sections

Tissues intended for cryosectioning were subjected to a cryoprotection procedure to minimize ice crystal formation during freezing. This involved sequential immersion in sucrose solutions of increasing concentration (10%, 20%, and 30% w/v). Cryosections were obtained using a sliding microtome equipped with a cooling stage, cutting the tissue at a thickness of 50 µm. The sections were collected in wells containing phosphate buffer (PB) and stored at 4 °C until use. Immunohistochemical labeling was performed on free-floating sections, with all incubations carried out in wells rather than on mounted slides. The immunostaining protocol followed was identical to that applied to paraffin-embedded material.

### Immunohistochemistry for Confocal Fluorescence Microscopy in Free-floating sections

A double immunofluorescence protocol was performed to carry out a detailed study of the mole brain. For this purpose, one of the two cerebral hemispheres from the specimen which had been fixed in formalin, was sectioned using a cryostat at a thickness of 50 μm. Three antibody pairs were used in the immunohistochemical analysis: OMP + MAP2, OMP + CR, and GAP43 + GFAP. Tissue sections were first washed three times for 5 minutes each in 0.1 M phosphate buffer (PB). Endogenous peroxidase activity was then blocked with 3% hydrogen peroxide (H₂O₂) to prevent background signal. Following an additional series of three 5-minute washes in PB, tissue permeability was enhanced by incubating the sections in 0.3% Triton X-100 for 15 minutes. After another three PB washes (5 min each), nonspecific binding sites were blocked using 2% bovine serum albumin (BSA) for 30 minutes.

Sections were then incubated with primary antibodies overnight at room temperature under gentle agitation. The following day, they were washed again (3 × 5 min in PB) and incubated with secondary antibodies for 30 minutes. Secondary antibodies were Alexa-conjugated fluorophores (NeoBiotech), specifically Alexa Fluor 488 anti-mouse IgG and Alexa Fluor 568 anti-rabbit IgG, both diluted 1:250 in PB. From this point onward, all steps were carried out in the dark to preserve fluorophore integrity. Sections were then washed in PB (3 × 10 min), incubated for 15 minutes in TO-PRO-3 (1:400 dilution), and finally rinsed again (3 × 5 min in PB) before mounting on glass slides. Imaging was performed using a Leica TCS SPE confocal system configured for dual-channel fluorescence detection.

### Image Acquisition

Digital micrographs were acquired using an Olympus SC180 camera mounted on an Olympus BX50 microscope (Tokyo, Japan). To obtain high-resolution images of extensive tissue regions, final composites were generated by merging up to 100 individual photomicrographs into seamless mosaics using PTGui automated stitching software (Rotterdam, Netherlands). Image processing was limited to uniform adjustments of brightness, contrast, and white balance, as well as cropping and standardization of image dimensions, all carried out in Adobe Photoshop CS4 (Adobe Systems, San Jose, CA). No further image modifications, enhancements, or artificial alterations were applied.

## Results

### Macroscopic study

A detailed macroscopic examination of the head and brain structures of the Iberian mole (*Talpa occidentalis*) revealed several anatomical landmarks relevant to the vomeronasal system and its associated pathways (Fig. 1). In the ventral view of the oral cavity (Fig. 1A), the incisive papilla was clearly identifiable along the midline of the hard palate, marking the external opening of the incisive duct and suggesting a functional route for the transmission of chemical cues to the vomeronasal organ (VNO). The dorsal view of the rostral nasal cavity (Fig. 1B) showed a pronounced skin folding, indicative of a highly specialized vestibular region adapted to the fossorial lifestyle. Rostral examination of the external nose (Fig. 1C) revealed a broad and flattened rhinarium, densely covered in vibrissae, consistent with a reliance on tactile and olfactory input in subterranean environments.

**Figure 1.**
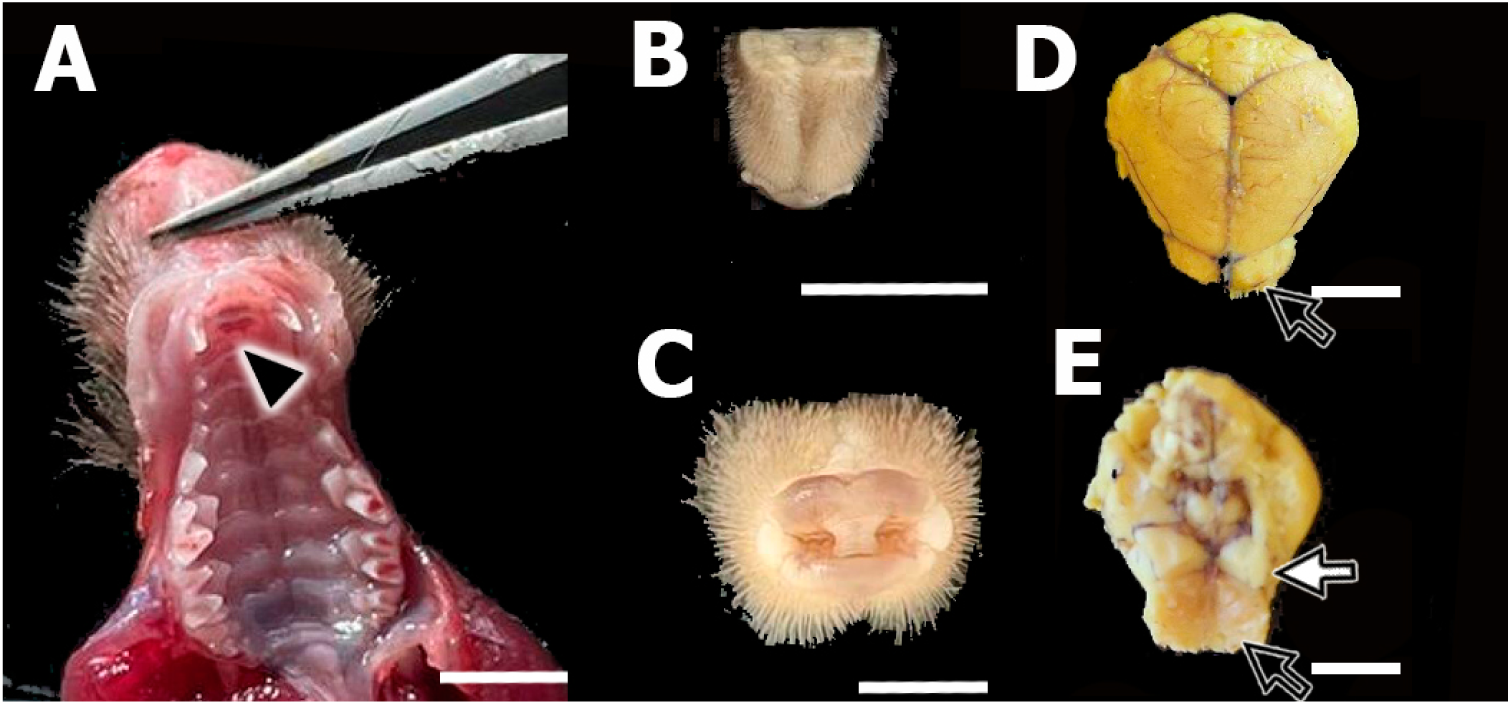
Macroscopic views of the anatomical regions under study in the Iberian mole (*Talpa occidentalis*). (A) Ventral view of the palatal mucosa, showing the location of the incisive papilla (arrowhead). (B) Dorsal view of the rostral region of the nasal cavity. (C) Rostral view of the rhinarium of *Talpa occidentalis*. (D) Dorsal view of the mole brain, where the olfactory bulbs (arrow) display a notably flattened morphology compared to other species. (E) Ventral view of the brain showing the concave ventral surface of the olfactory bulbs (black arrow) and the olfactory tubercules (white arrow). Scale bars: 1 cm (A, B, D, E); 0.5 cm (C).

In the dorsal view of the encephalon (Fig. 1D), the olfactory bulbs appeared notably flattened and compressed dorsoventrally compared to other small mammals, with reduced lateral expansion. This compact morphology may reflect ecological constraints associated with fossorial adaptation and cranial space limitation. The ventral aspect of the brain (Fig. 1E) further illustrated the concave ventral surface of the olfactory bulbs, which were separated by a shallow interbulbar groove. Posteriorly, the olfactory tracts projected symmetrically. Notably, the olfactory tubercle appeared conspicuously developed, occupying a substantial portion of the ventral forebrain and suggesting an important integrative role in olfactory processing in this species.

### Histological and immunohistochemical study of the central VNO

In the central region of the nasal cavity of *Talpa occidentalis*, transverse sections revealed the anatomical configuration of the VNO in close relation to adjacent structures (Fig. 2). The VNO was positioned ventrally to the dorsal and ventral nasal turbinates and dorsally to the palatal mucosa, enclosed by the vomeronasal cartilage and flanked medially by the nasal septum and ventrally by the vomer bone (Fig. 2C). The nasolacrimal duct was identified laterally (Fig. 2C). One of the specimens analyzed presented signs of vomeronasalitis, with prominent inflammation localized to the lateral portion of the left vomeronasal duct (VD). This was evidenced by epithelial disruption and the presence of numerous inflammatory cells within the lumen and subepithelial region (Fig. 2A–B).

**Figure 2.**
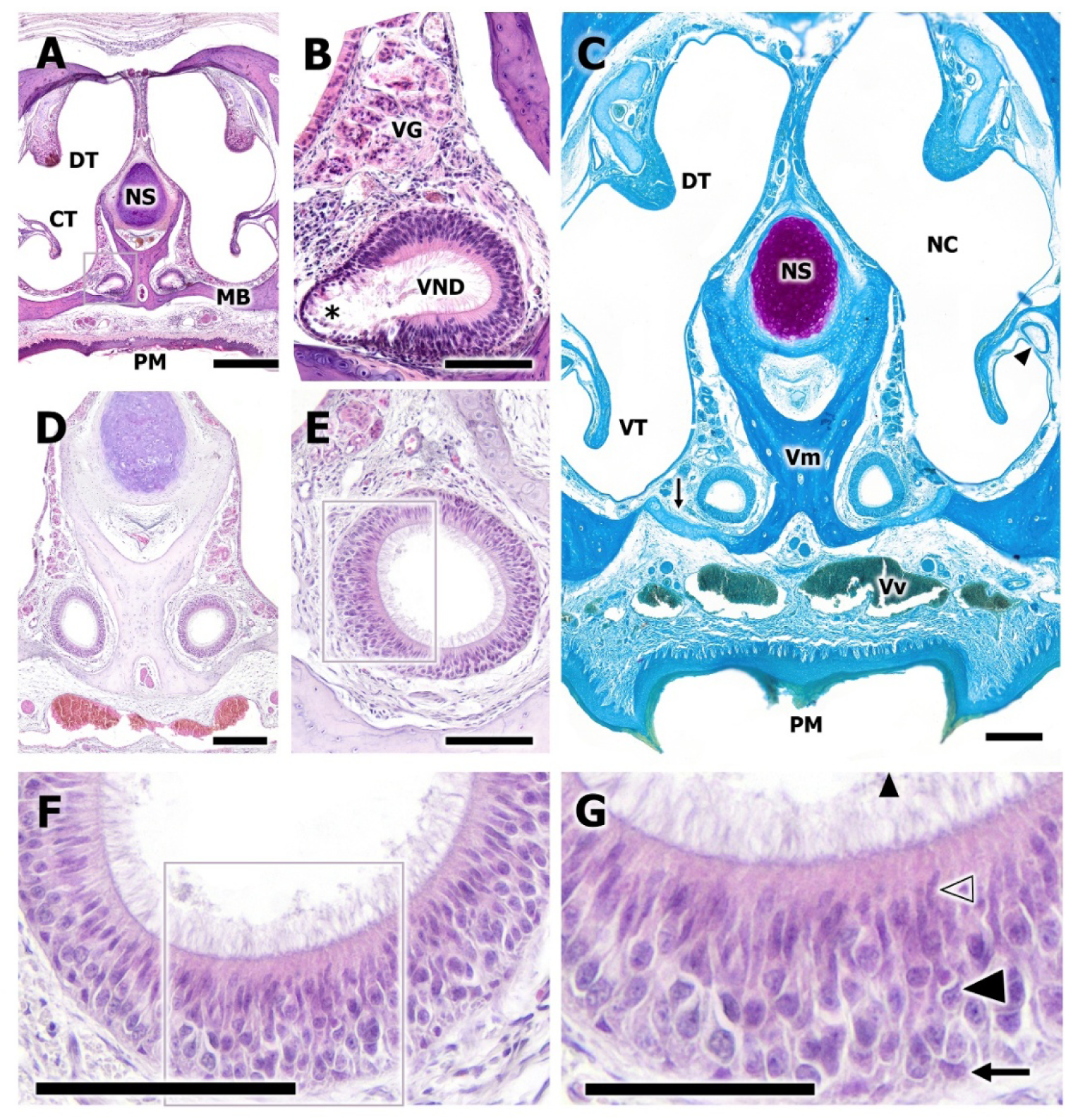
Microscopic study of the central region of the VNO in *Talpa occidentalis*. (A– B) Transverse overview of the nasal cavity (NC) of the mole. The specimen analyzed exhibited vomeronasalitis affecting the lateral portion of the vomeronasal duct. The enlargement of the left VNO reveals epithelial damage due to inflammation and a high concentration of inflammatory cells (). (C) Transverse view of the NC showing the topographical relationships of the VNO with the vomer bone (Vm), nasal septum (Ns), palatal mucosa (Pm), and vomeronasal cartilage (arrow). Also visible are the dorsal (Dt) and ventral (Vt) nasal turbinates and the nasolacrimal duct (arrowhead). (D) The VNOs exhibit a close anatomical relationship with the vomer bone, adapting precisely to its lateral curvatures. The vomeronasal glands (VG) are highly developed and located dorsolaterally to the vomeronasal duct. (E) Enlargement of the left VNO duct from D. The duct has a circular cross-section, lined by a continuous SE of substantial thickness and regular morphology, without lateral-medial differentiation. Gallego’s Trichrome stain. (F) Higher magnification of the SE from the previous image. The pronounced development of apical ciliary projections into the lumen is noteworthy. (G) Enlargement of the epithelial layers shown in image F. The pseudostratified epithelium is composed of basal cells (arrow), bipolar sensory neurons (arrowhead), sustentacular cells (open arrowhead), and cilia (). All sections were stained with H&E, except for image E, which used Gallego’s Trichrome. Scale bars: 500 μm (A), 200 μm (C, D), 100 μm (B, D, F, G).

In well-preserved regions, the VNO duct showed a circular cross-section with a continuous and thick layer of sensory epithelium (SE) exhibiting no clear lateral-medial zonation (Fig. 2E). At higher magnification, this epithelium presented well-organized apical ciliary projections extending into the lumen (Fig. 2F), and a pseudostratified cellular organization composed of basal progenitor cells, bipolar sensory neurons, sustentacular cells, and dense tufts of cilia (Fig. 2G). Moreover, the vomeronasal glands appeared well-developed, in a dorsolateral position with respect to the duct, and conforming closely to the curvature of the vomer bone (Fig. 2D).

Histochemical staining with Alcian Blue and PAS revealed specific features of the mucosal environment surrounding the VNO (Fig. 3). Alcian Blue staining, which highlights acidic mucopolysaccharides, showed prominent affinity in the vomeronasal septum and the inner lining of the dorsal nasal turbinates, as seen in the overview of the nasal cavity (Fig. 3A). In the mid-regional section of the nasal cavity, slight staining was also detected in the vomeronasal cartilage (Fig. 3C), whereas in more caudal sections (Fig. 3D), positive Alcian Blue staining was apparent in the respiratory mucosa and in the lumen of both vomeronasal ducts, indicating the presence of acidic secretions in both regions.

**Figure 3.**
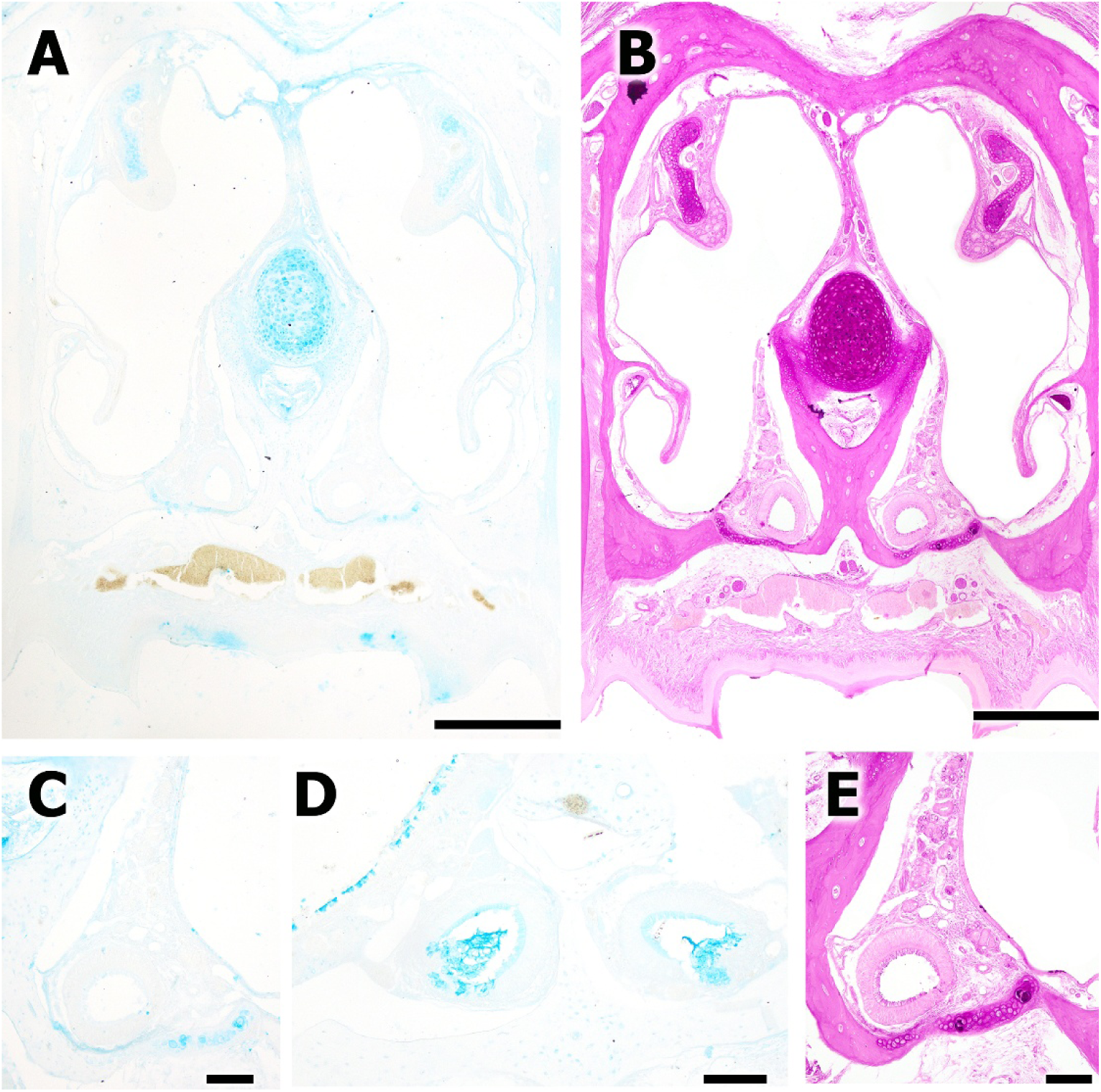
Histological study of the VNO in *Talpa occidentalis* using Alcian Blue (AB) and Periodic Acid-Schiff (PAS) staining. A, C, D. VNO stained with Alcian Blue. A. General view of the nasal cavity showing intense staining of the vomeronasal septum and the inner surface of the dorsal turbinates. C. Mid-level view of the VNO. Slight Alcian Blue staining is observed in the vomeronasal cartilage. D. Caudal view of both vomeronasal ducts. Positive AB staining is evident in the respiratory mucosa and within the lumen of the VNO. B, E. PAS staining. B. Global view of the nasal cavity at the central level of the VNO, showing general staining of all structures. E. Higher magnification of the vomeronasal duct from image B. Intense staining is observed in the apical region of the sensory epithelium and within the glandular plexus. Scale bars: 500 μm (A, B), 100 μm (C, D, E).

Periodic Acid-Schiff (PAS) staining, which detects neutral glycoconjugates, revealed a broad labeling pattern across all structures at the mid-level of the nasal cavity (Fig. 3B). At higher magnification, the apical region of the sensory epithelium and the associated glandular plexus showed intense PAS reactivity (Fig. 3E), suggesting a strong presence of neutral mucosubstances in both the epithelial and subepithelial compartments of the VNO.

The immunohistochemical study of the VNO at its central level was performed to explore the molecular organization of its sensory and associated structures (Fig. 4 and 5). Immunolabeling for the Gαo subunit showed positive staining in both VNOs, located symmetrically on either side of the vomer bone (Fig. 4A). The signal was not restricted to the sensory epithelium, as partial immunopositivity was also detected in the vomeronasal glands and associated nerve bundles. Higher magnification of the left duct (Fig. 4B) revealed clear immunoreactivity in the sensory epithelium, with an especially strong signal in the apical region.

**Figure 4.**
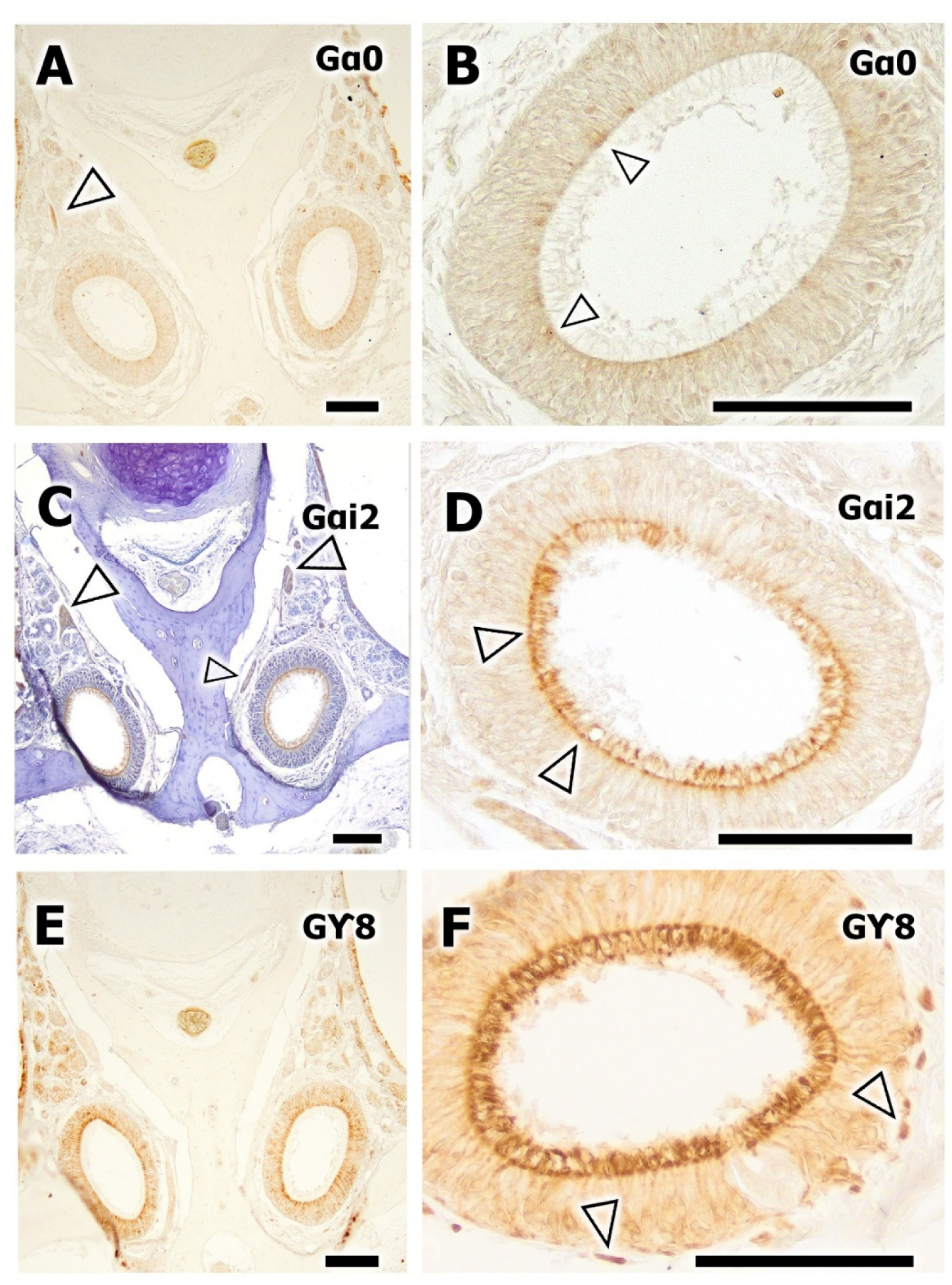
Immunohistochemical study of G protein subunits in the mole VNO at the central level. A–B. Immunohistochemical labeling for Gαo. A. General view of both VNOs located on either side of the vomer bone. Immunopositivity is observed in both VNOs and partially in the vomeronasal glands and nerves (open arrowhead). B. Enlargement of the left duct from image A. Immunoreactivity is evident in the SE, especially in its apical zone (open arrowhead). C–D. Immunolabeling for Gαi2. C. General view of both vomeronasal ducts within their anatomical context. Strong immunostaining for the Gαi2 subunit is observed in the dorsal and medial vomeronasal nerves (open arrowhead). Hematoxylin counterstaining. D. Enlargement of the right vomeronasal duct from image C. Intense immunolabeling for Gαi2 is visible in the cilia and dendritic knobs of sensory neurons in the vomeronasal epithelium (open arrowhead), as well as in vomeronasal axons. E–F. Immunohistochemical labeling for Gγ8. E. General view of both vomeronasal ducts and the surrounding vomeronasal parenchyma. Positive immunolabeling is observed in vomeronasal glands. F. Central vomeronasal duct showing strong immunoreactivity in cilia, the apical portion of vomeronasal neurons, and their axons in the lamina propria (open arrowhead). Scale bars: 100 μm.

**Figure 5.**
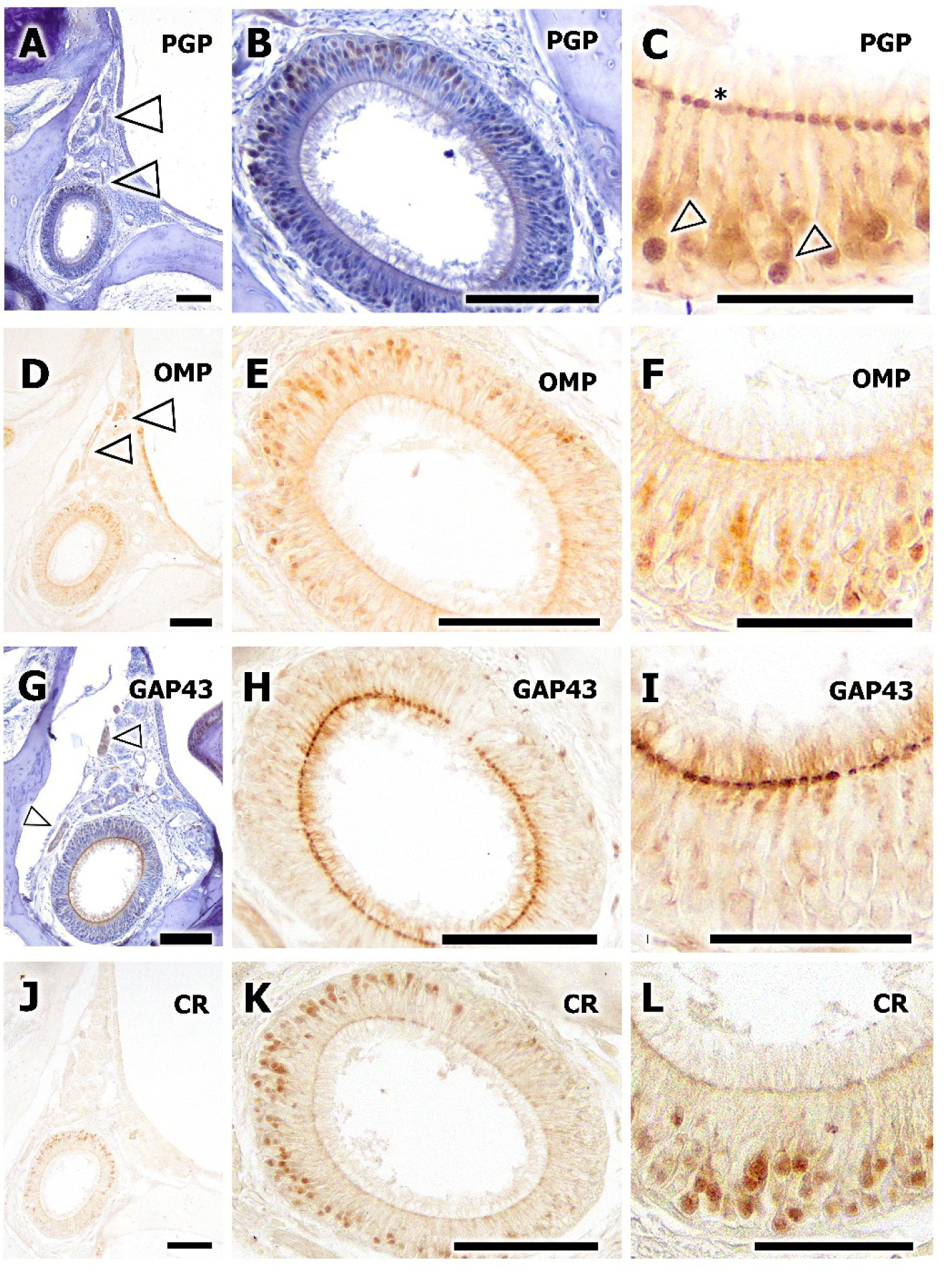
Immunohistochemical study of the central VNO using anti-PGP, -OMP, - GAP43, and -CR markers. A–C. Anti-PGP. A–B. Hematoxylin counterstaining. A. Transverse section of the VNO. Immunoreactivity is observed in dorsomedial bundles of vomeronasal nerves (arrowhead). B. Enlargement of the VD. Strong labeling of the cell bodies of vomeronasal bipolar neurons, especially in the dorsal SE. C. Enlargement of the dorsal SE from the previous image. Intense expression of PGP is seen in neuronal somata (open arrowhead) and in the apical knobs of bipolar neurons. Weaker labeling is noted in the dendritic projections extending into the lumen. D–F. Anti-OMP. D. Transverse section of the VD and surrounding parenchyma. Negative labeling for OMP throughout the parenchyma, except in some dorsal nerve bundles (open arrowhead). E. Enlargement of the VD from image D. Positive labeling in the apical area of the SE and in the neuronal cell bodies of bipolar neurons in the dorsal half. F. Enlargement of the dorsal SE from E. Anti-OMP primarily labels cell bodies of vomeronasal bipolar neurons. G–I. Anti-GAP43. G. Immunostaining in the parenchyma is limited to dorsal bundles of vomeronasal nerves (open arrowhead). H. Enlargement of the VD from image G. Strong immunoreactivity is observed in the apical knobs of bipolar neurons throughout the sensory epithelium. I. Enlargement of the SE from image H. Anti-GAP43 antibody labels the apical knobs and their cilia. J–L. Anti-CR antibody. J. No CR immunoreactivity is detected in the vomeronasal parenchyma. K–L. Enlargement of the VD from image J (K) and of the epithelium from image K (L). The labeling is concentrated in the cell bodies of vomeronasal bipolar neurons. Scale bars: 100 μm (A, B, D, E, G, H, J, K), 50 μm (C, F, I, L).

Labeling for the Gαi2 subunit (Fig. 4C–D) showed a complementary pattern. At low magnification (Fig. 4C), the dorsal and medial vomeronasal nerves displayed intense immunoreactivity. The epithelium itself also showed a distinct labeling pattern, as confirmed in the enlargement of the right VNO duct (Fig. 4D), where Gαi2 was concentrated in the dendritic knobs and cilia of the sensory neurons, as well as in their axons projecting into the lamina propria.

The expression of the Gγ8 subunit (Fig. 4E,F) was particularly evident in the vomeronasal glands (Fig. 4E). Additionally, intense staining was observed in the cilia, the apical region of sensory neurons, and their axons within the lamina propria of the central VNO duct (Fig. 4F).

Immunolabeling with the anti-PGP antibody revealed strong immunoreactivity in dorsomedial vomeronasal nerve bundles (Fig. 5A). The sensory epithelium, especially its dorsal portion, showed intense staining of the cell bodies of bipolar vomeronasal neurons (Fig. 5B). A detailed examination of this region (Fig. 5C) showed that PGP expression was localized both to the neuronal somata and to the apical knobs of sensory neurons, with lighter staining extending into their dendritic projections within the lumen. The anti-OMP antibody showed a different pattern. In transverse sections, most of the vomeronasal parenchyma appeared negative, except for certain dorsal nerve bundles that retained some immunoreactivity (Fig. 5D). However, in the dorsal half of the sensory epithelium, OMP labeling was clearly present in the apical domain and in the neuronal somata of bipolar neurons (Fig. 5E–F).

Immunolabeling with the anti-GAP43 antibody was strongly restricted to neuronal elements. The parenchyma only exhibited signal in dorsal vomeronasal nerve bundles (Fig. 5G), but in the sensory epithelium (Fig. 5H), robust labeling was evident in the apical knobs of bipolar neurons across the entire epithelium. At higher magnification, this marker also stained the cilia projecting from these knobs into the ductal lumen (Fig. 5I). The anti-CR antibody, in contrast, did not produce detectable immunoreactivity in the general parenchyma (Fig. 5J). Nevertheless, more detailed views of the vomeronasal duct (Fig. 5K–L) demonstrated CR expression in the cell bodies of some bipolar neurons.

Histochemical analysis of the central VNO in *Talpa occidentalis* using lectins LEA, SBA, ECL, and WGA revealed distinct glycoconjugate distribution patterns in both the sensory epithelium and associated tissues (Figs. 6 and 7). LEA labeling showed strong affinity for the vomeronasal glands and for the mucosal lining of the ventral turbinates (Fig. 6A). The sensory epithelium displayed intense LEA binding, particularly in the cilia, apical dendritic knobs, and basal sensory receptor cells (Fig. 6B). Staining with SBA demonstrated a more selective distribution. In transverse sections, positive SBA reactivity was detected in the vomeronasal cartilage and in a subset of vomeronasal glands (Fig. 6C), with additional strong labeling in the nasolacrimal duct. In the sensory epithelium, SBA marked the apical surface and cilia of bipolar neurons (Fig. 6F).

**Figure 6.**
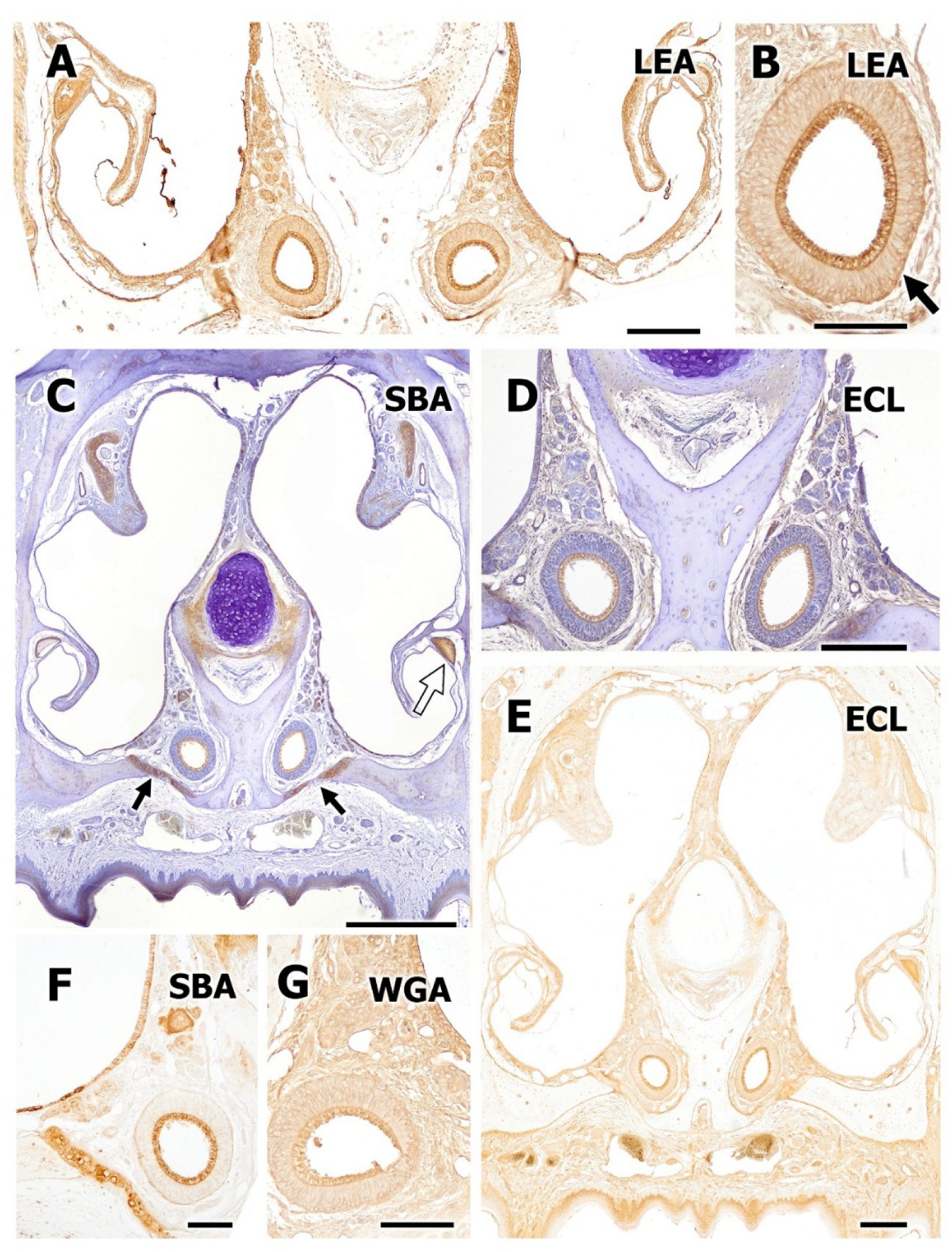
Histochemical study of the central VNO of the Iberian mole using the lectins LEA, SBA, ECL, and WGA. A–B. Histochemical labeling with LEA. A. General view of the vomeronasal organs, vomer bone, and ventral turbinates, showing intense labeling in vomeronasal parenchymal glands and in the respiratory and lining mucosa of the ventral turbinates. B. Enlargement of the left vomeronasal duct from image A. Positive histochemical reaction highlighting the cilia of the sensory epithelium, apical dendritic knobs, and sensory receptor cells, particularly those located in the basal region (arrow). C, F. Histochemical labeling with SBA. C. Transverse section of the nasal cavity showing positive SBA staining in the vomeronasal cartilage and in a subpopulation of vomeronasal glands. Strong positivity is also noted in the nasolacrimal duct (open arrow). F. Enlargement of the left vomeronasal organ from image C. Positive SBA staining is observed in the cilia of the vomeronasal sensory epithelium and on the apical surface of the neuroepithelium. D–E. Histochemistry with ECL lectin. D. Enlarged view of both VNOs from image E, counterstained. Intense positive labeling is observed in the cilia of the sensory epithelium. E. General view of the nasal cavity showing moderately positive staining throughout, with stronger intensity on the mucociliary surface of the vomeronasal duct. G. Histochemical staining with WGA. Diffuse positive labeling throughout the parenchyma and duct, with stronger signal in the cilia. Hematoxylin counterstaining in C and D. Scale bars: 500 μm (C), 200 μm (A, D, E), 100 μm (B, F, G).

**Figure 7.**
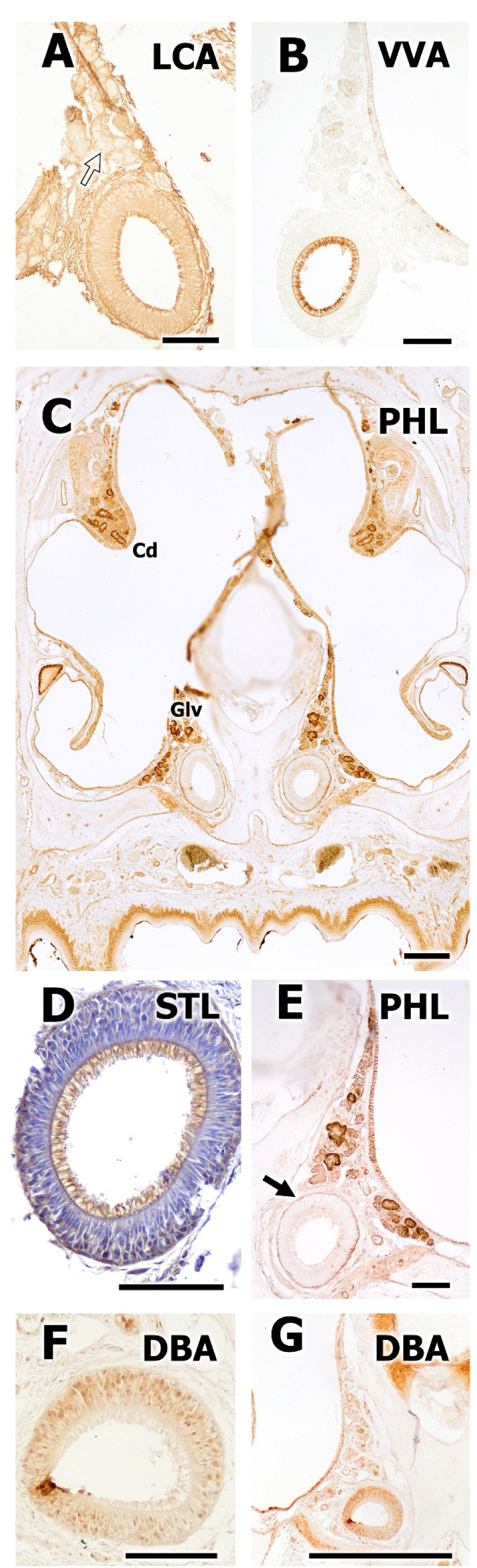
Histochemical study of the central vomeronasal organ of the Iberian mole using the lectins LCA, VVA, PHL, STL, and DBA. A. LCA histochemical labeling shows positive staining throughout the vomeronasal duct, with stronger intensity in the apical and basal regions of the sensory epithelium. In the vomeronasal parenchyma, staining is also positive, except in the vomeronasal glands, which appear negative (open arrow). B. VVA labeling is highly intense and specific in the cilia and apical knobs of the sensory epithelium. C, E. PHL histochemistry. C. General view of the nasal cavity, with strong staining in the glands of the vomeronasal parenchyma (Gv) and the dorsal turbinates (Cd). E. Enlargement of the right VNO from image C shows intense staining in the basal cell band of the sensory epithelium. D. STL histochemistry with H&E counterstaining shows strong labeling in the cilia and basal cells of the vomeronasal sensory epithelium. F–G. DBA lectin staining. G. General view of the VNO with intense labeling in the cartilage and VD, and weaker labeling in the vomeronasal glands. F. Enlargement of the vomeronasal duct from image G reveals staining in the cell bodies of bipolar neurons in the vomeronasal sensory epithelium. Scale bars: 500 μm (G), 100 μm (A-F).

**Figure 8.**
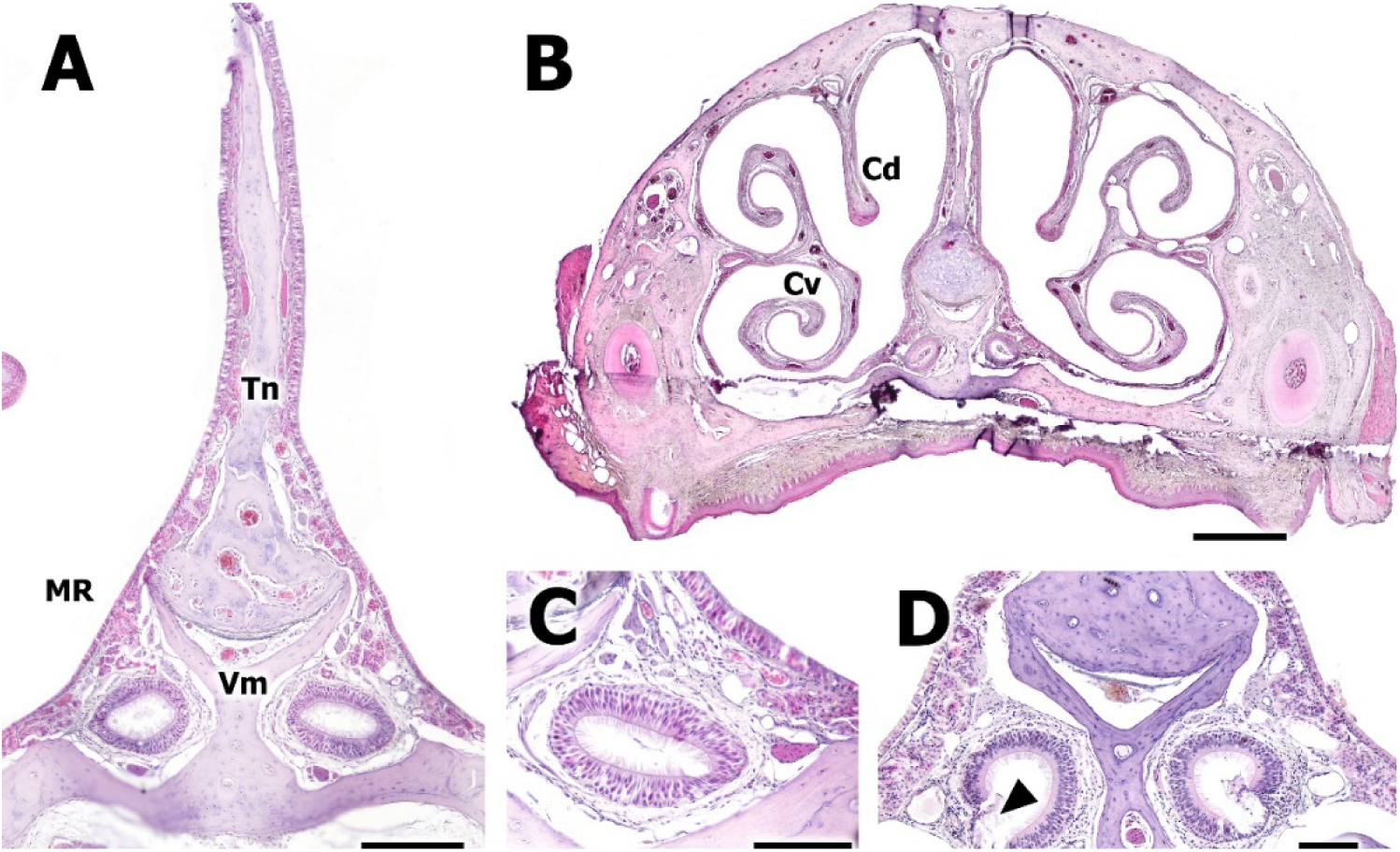
Microscopic analysis of the vomeronasal organ of *Talpa occidentalis* at a caudal level. A. View of both vomeronasal ducts. A global reduction of the vomeronasal parenchyma is observed, along with a ventromedial displacement of the ducts caused by a decrease in the size of the vomer bone (Vm). B. General view of the caudal nasal cavity of the mole. A marked development of the dorsal and ventral nasal conchae (Cd, Cv) is noted, occupying a large portion of the nasal cavity volume. C. At higher magnification, the VNO loses the rounded shape characteristic of the central level. As the duct progresses caudally, it contracts dorsoventrally until it eventually reduces and disappears. However, epithelial features are preserved throughout its perimeter, with identifiable cell types and surface cilia. D. View of two caudal vomeronasal organs affected by vomeronasalitis. Despite retaining a morphology similar to the central duct, sensory epithelial degradation is observed (arrowhead), indicating the presence of vomeronasalitis. Nevertheless, the dorsomedial and ventral portions of the duct maintain functionality. Scale bars: 500 μm (B), 200 μm (A), 100 μm (C,D).

ECL labeling revealed a more epithelial-specific pattern. The cilia of the sensory epithelium showed strong staining (Fig. 6D), while the entire nasal cavity displayed a moderately positive signal, with the most intense labeling observed on the mucociliary surface of the vomeronasal duct (Fig. 6E). Finally, WGA staining showed a diffuse but consistent pattern throughout the vomeronasal duct and surrounding parenchyma (Fig. 6G), with enhanced intensity in the cilia. This pattern suggests widespread expression of WGA-binding glycoconjugates in both epithelial and subepithelial compartments.

LCA staining was widespread throughout the VD, with stronger labeling localized to both the apical and basal areas of the sensory epithelium (Fig. 7A). The surrounding parenchyma also showed LCA reactivity, though the vomeronasal glands remained unstained. Labeling with VVA showed a highly specific and intense signal in the apical knobs and cilia of the sensory epithelium (Fig. 7B), suggesting selective glycosylation features in the apical domain of sensory neurons.

PHL labeling revealed a clear preference for glandular structures, bound intensely to glands in both the vomeronasal parenchyma and the dorsal turbinates (Fig. 7C). It also stained the basal layer of cells within the sensory epithelium (Fig. 7E). STL histochemistry, with hematoxylin counterstaining, showed a distinct pattern of reactivity in the cilia and in the basal cells of the sensory epithelium (Fig. 7D), further supporting the presence of multiple glycosylation compartments within the neuroepithelium. Finally, DBA labeling revealed a dual pattern, staining both the vomeronasal duct and the surrounding cartilage intensely, while showing weaker labeling in the vomeronasal glands (Fig. 7G). Upon closer examination, the cell bodies of bipolar neurons within the sensory epithelium were also positively labeled (Fig. 7F), suggesting specific carbohydrate moieties associated with neuronal membranes.

### Histological and immunohistochemical study of the caudal VNO

At the caudal level of the VNO, several anatomical and histological changes were observed (Fig. 11). A noticeable reduction in the overall vomeronasal parenchyma was accompanied by a ventromedial displacement of the ducts (Fig. 11A), likely resulting from a decrease in the size of the vomer bone. In this same region, the nasal cavity showed substantial development of the dorsal and ventral conchae (Fig. 11B), which occupied a considerable portion of the available space. As the vomeronasal duct advanced caudally, it progressively lost the rounded profile typical of the central level, exhibiting a marked dorsoventral contraction until it diminished and eventually disappeared (Fig. 11C). Despite this morphological regression, the epithelium maintained its characteristic features, including the presence of identifiable cell types and surface cilia along the luminal perimeter. In some specimens, caudal segments of the VNO showed evidence of vomeronasalitis (Fig. 11D): although the general shape of the duct remained similar to that of the central level, partial degradation of the sensory epithelium was evident. Nonetheless, the dorsomedial and ventral portions of the duct retained structural integrity, suggesting the persistence of functional capacity in these regions.

The caudal VNO exhibited distinct patterns of immunoreactivity when analyzed using antibodies against CR, PGP, GAP43, and Gα0 (Fig. 9). Immunostaining for calretinin (CR) revealed intense labeling in the dorsal vomeronasal nerves (Fig. 9A), while a weak signal was present in restricted regions of the sensory epithelium, particularly in the apical portions of bipolar vomeronasal neurons (Fig. 9A). The neuronal marker PGP produced strong immunoreactivity in vomeronasal nerves throughout the caudal VNO (Fig. 9B), including prominent staining of the dorsal nerve branches (Fig. 9C). At higher magnification, PGP labeling was also evident in the apical zone of bipolar neurons (Fig. 9D), as well as in ventral vomeronasal nerve fibers (Fig. 9D). GAP43 immunostaining further confirmed strong labeling of vomeronasal nerves lateral to the vomer (Fig. 9E), and marked expression in the apical region of bipolar neurons within the epithelium (Fig. 9F). Lastly, immunoreactivity for the Gα0 protein subunit was observed in vomeronasal nerves running along the nasal septum (Fig. 9G), and in the sensory epithelium of the vomeronasal duct, with staining present in both apical neuronal compartments and associated axons (Fig. 9H).

**Figure 9.**
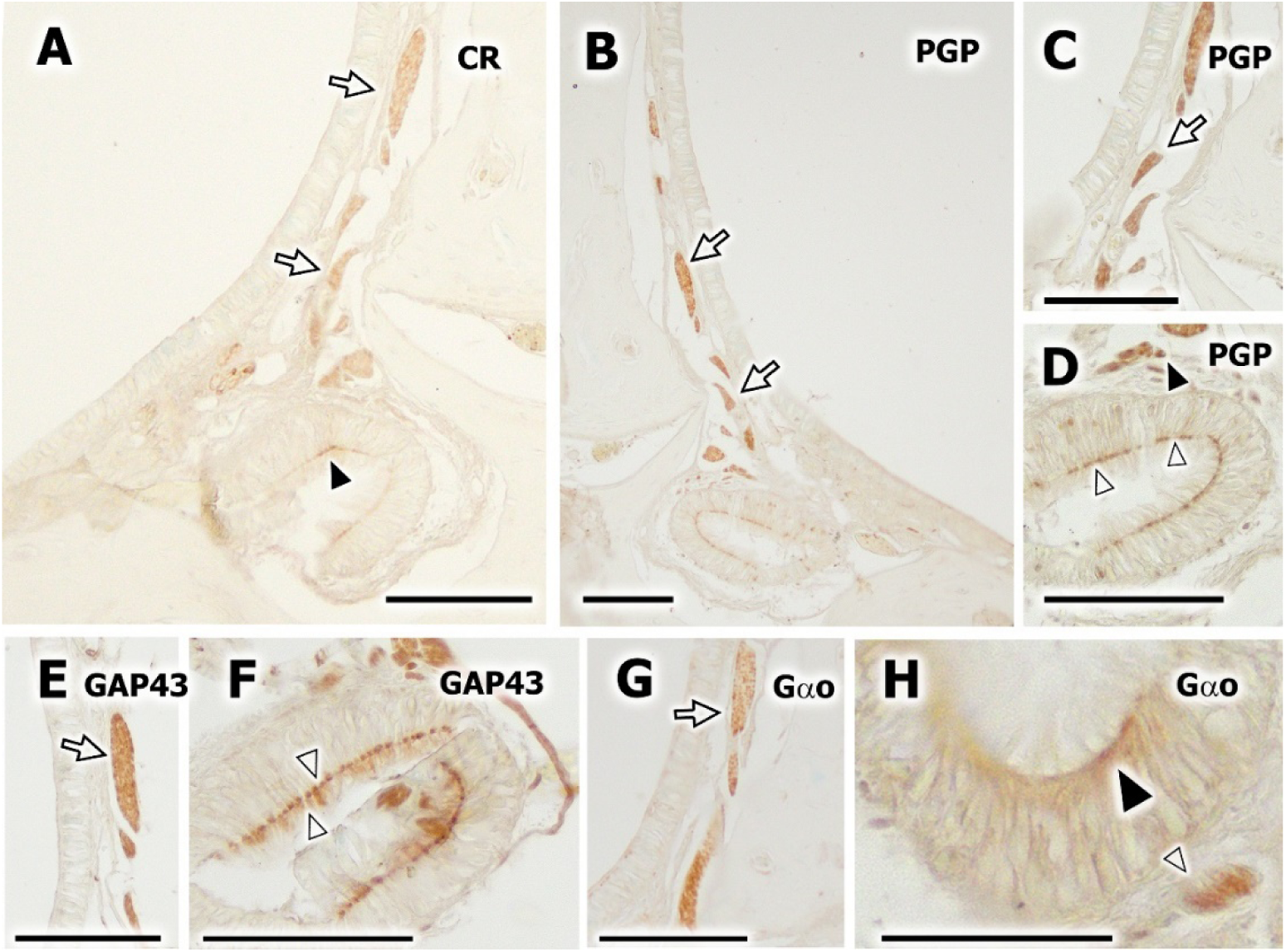
Immunohistochemical analysis of the caudal vomeronasal organ using antibodies against CR, PGP, GAP43, and Gα0. A. Immunostaining for the calcium-binding protein CR. An intense signal is observed in the dorsal vomeronasal nerves (open arrow), and a weak signal is detected in specific regions of the vomeronasal SE (arrowhead), particularly in the apical zone of bipolar vomeronasal neurons. B–D. Immunohistochemical labeling for the neuronal marker PGP. B. General view of the caudal VNO, showing strong staining of the vomeronasal nerves (open arrow). C. Higher magnification of the dorsal vomeronasal nerves, which are intensely labeled by the anti-PGP antibody (open arrow). D. High magnification of the VD from image B, showing intense PGP staining in the apical region of bipolar vomeronasal neurons (open arrowhead) and in vomeronasal nerves located ventrally to the VNO (arrowhead). E–F. Immunostaining for the protein GAP43. E. Transverse section of vomeronasal nerves (open arrow), located laterally to the vomer bone, showing strong positive staining for the anti-GAP43 antibody. F. Vomeronasal duct displaying intense apical staining in bipolar neurons of the sensory epithelium (open arrowhead). G–H. Immunolabeling for the Gα0 subunit. G. Section of vomeronasal nerves running through the nasal septum (open arrow), which show intense Gα0 immunoreactivity. H. Sensory epithelium of the caudal VD with positive staining in the apical region (arrowhead) and in vomeronasal axons (open arrowhead). Scale bar: 100 μm.

Lectin histochemistry performed on the caudal VNO revealed differential binding patterns depending on the carbohydrate affinity of each probe (Fig. 10). VVA staining showed scattered positive cells across the vomeronasal sensory epithelium (Fig. 10A), with a similar pattern observed in the adjacent respiratory mucosa (Fig. 10A). A comparable distribution was found with DBA, which labeled isolated cells in both the vomeronasal epithelium and the respiratory lining (Fig. 10B). STL exhibited a more generalized pattern, strongly marking the entire epithelium of both respiratory and vomeronasal origins (Fig. 10C). SBA lectin showed diffuse reactivity within the vomeronasal epithelium (Fig. 10D), as well as in the respiratory mucosa (Fig. 10D). ECL histochemistry revealed a uniform and intense staining across the entire extent of the caudal VNO (Fig. 10E), while PHL also demonstrated a global labeling pattern affecting both the vomeronasal and respiratory mucosae (Fig. 10F).

**Figure 10.**
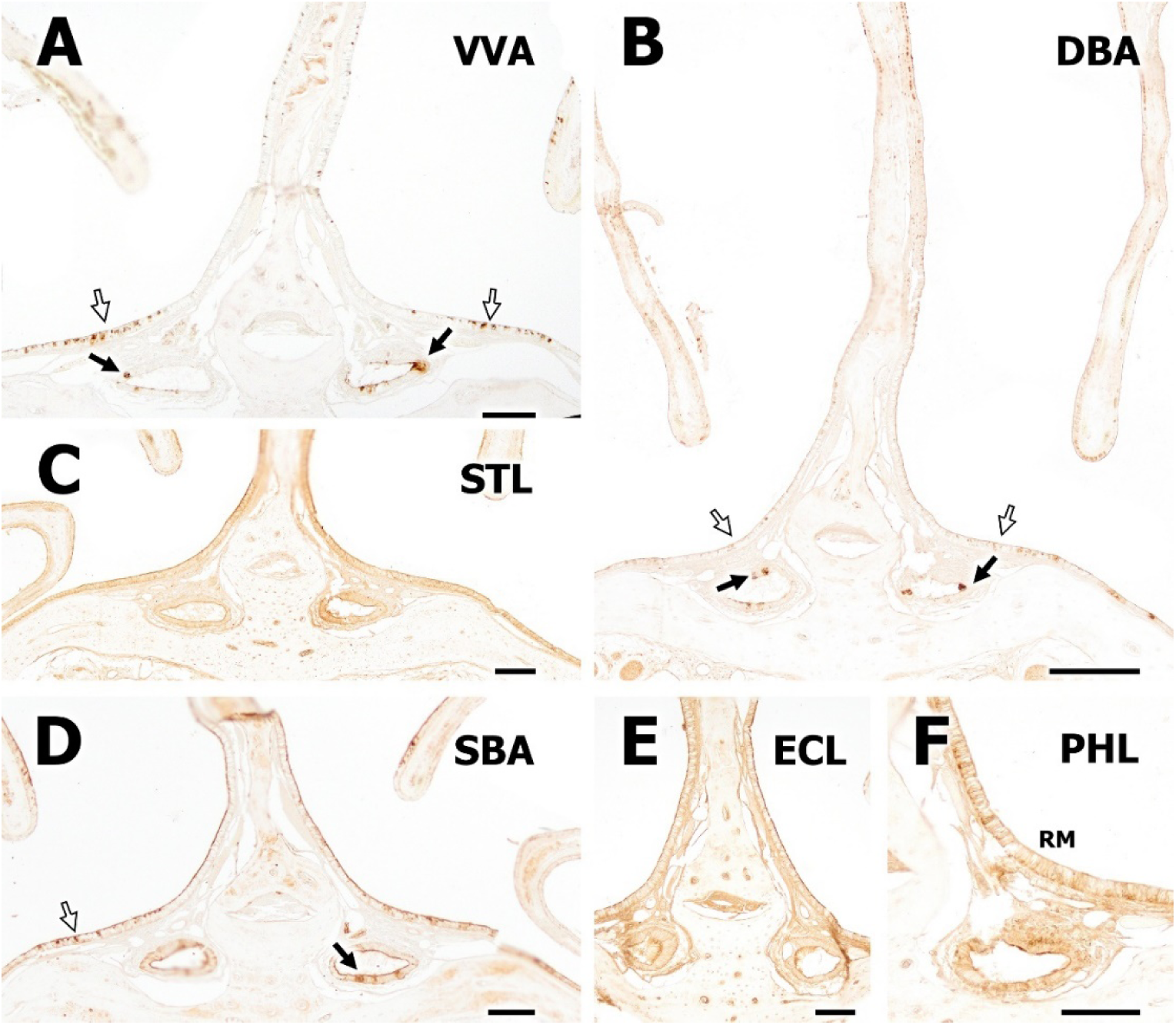
Histochemical analysis with the lectins VVA, DBA, STL, SBA, ECL, and PHL in the caudal vomeronasal organ. A. VVA histochemistry. General view of the two caudal VNOs showing a positive pattern in scattered cells of the SE (arrows). The respiratory mucosa exhibits a similar labeling pattern (open arrows). B. DBA labeling. General view of the VDs, showing positive staining in isolated sensory epithelial cells (arrows) and in the respiratory mucosa (open arrows). C. STL histochemistry in the caudal VNO. This lectin displays a generalized pattern throughout both the respiratory and vomeronasal sensory epithelium. D. SBA labeling shows diffuse staining in the vomeronasal sensory epithelium (arrow) and in the respiratory mucosa (open arrow). E. ECL histochemistry in the caudal VNO reveals a uniformly positive pattern throughout the structure. F. PHL labeling shows a global staining pattern of both the respiratory mucosa (RM) and the vomeronasal epithelium. Scale bars: 200 μm (B), 100 μm (A, C, D, E, F).

### Microscopic Study of the Incisive Papilla

The incisive papilla of exhibited distinct structural and topographic features upon microscopic examination (Fig. 11). A general view revealed the papilla positioned between and ventral to the incisor teeth, which surrounded it dorsally and bilaterally (Fig. 11A). At higher magnification, the papillary surface showed a pronounced interdigitation between the connective tissue papillae of the lamina propria and the invaginations of the overlying epithelium (Fig. 11B).

**Figure 11.**
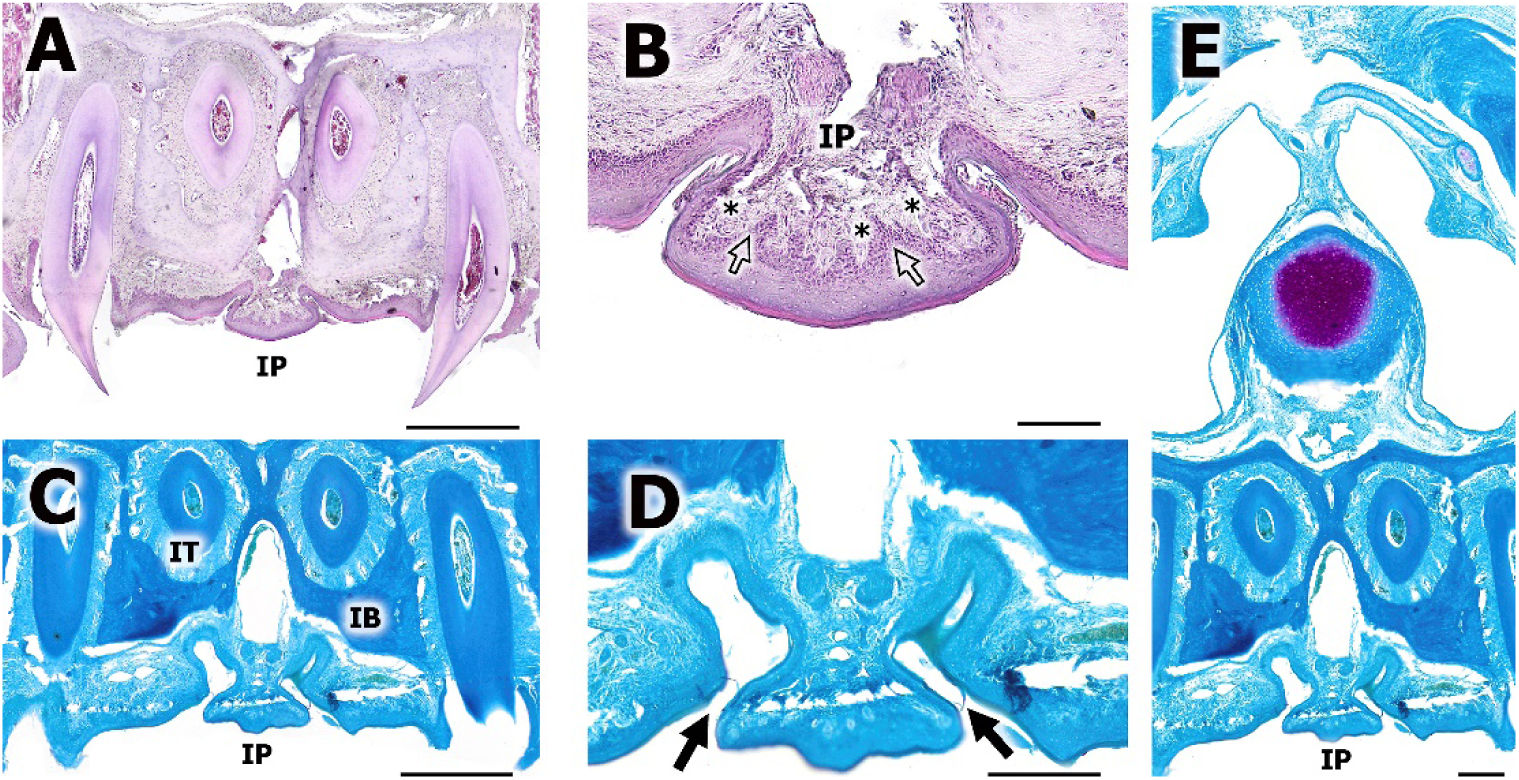
Histological study of the incisive papilla of *Talpa occidentalis*. A. General view of the papilla showing its topographic relationships. The incisor teeth (IT) are observed surrounding the papilla dorsally and laterally. B. Higher magnification of the papilla from image A, showing the marked interdigitation between the connective tissue papillae of the palatal lamina propria (*) and the epithelial invaginations of the papillary surface. C–E. Study of the incisive papilla using Gallego’s trichrome staining. C. Enlarged view of image E, displaying the papilla, the roots of the incisor teeth (IT), and the incisive bone (IB). D. Higher magnification of image C, where the papilla and the initial formation of the incisive ducts flanking it can be seen (arrows). E. General overview of the papilla in relation to the structures of the nasal septum. Histological stains: HE (A, B), Gallego’s trichrome (C, D, E). Scale bars: 500 μm (A, C), 200 μm (D, E), 100 μm (B).

Gallego’s trichrome staining further clarified the relationships between the papilla and surrounding skeletal structures. A detailed view demonstrated the proximity of the papilla to the roots of the incisor teeth and to the incisive bone (Fig. 11C), while a subsequent magnification revealed the early formation of the incisive ducts flanking the papilla on both sides (Fig. 11D). Finally, a broader histological overview allowed the visualization of the papilla in the context of the nasal septum and adjacent anatomical elements (Fig. 11E).

The communication between the vomeronasal and incisive ducts was additionally examined through serial transverse histological sections stained with hematoxylin-eosin (Fig. 12). In the most rostral sections, the ducts appeared as separate structures: the incisive ducts were located ventrally and exhibited a narrow, elongated morphology, whereas the vomeronasal ducts were more dorsal and oval in shape (Fig. 12A). Progressing caudally, the two ducts were observed to converge and establish a communication, ultimately merging into a single passage (Fig. 12B). At this level, the palatal mucosa also contained the trigeminal nerve and a prominent glandular plexus (Fig. 12B). A closer view confirmed the anatomical connection between the two ducts, with the vomeronasal and incisive lumina visibly merging (Fig. 12C). Additional sections showed the trajectory of the incisive ducts ascending dorsally toward the vomeronasal ducts (Fig. 12D). In more caudal regions, a lateral displacement of the incisive ducts was noted, positioning them beside the vomeronasal ducts and aligning with the palatal fissure (Fig. 12E). A final overview contextualized the relationship of both duct systems within the nasal cavity, emphasizing their spatial arrangement and association with the vomeronasal cartilage (Fig. 12F).

**Figure 12.**
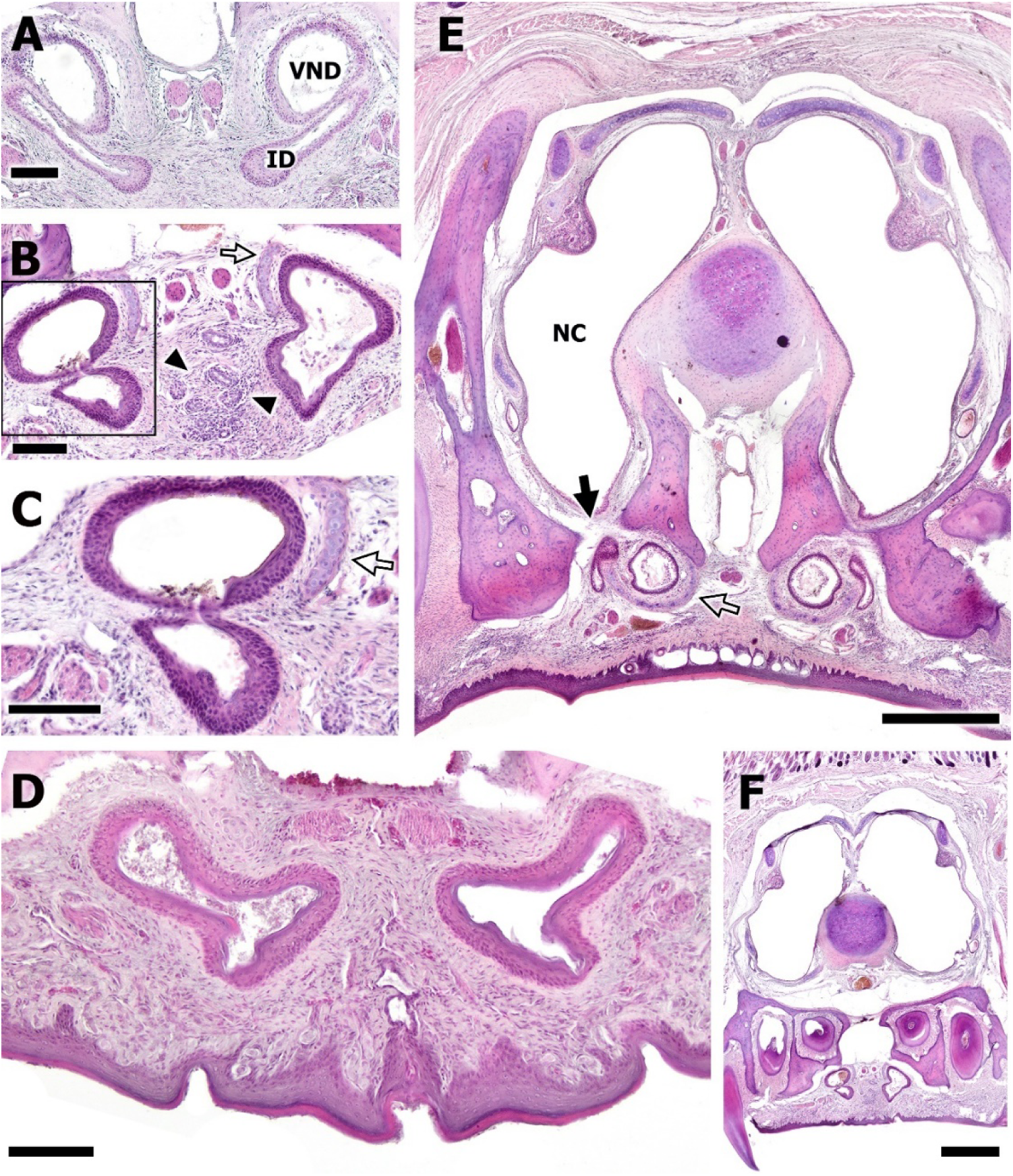
Serial microscopic analysis in the transverse plane of the communication between the vomeronasal and incisive ducts using hematoxylin-eosin staining. A. Section of the ventral region of the nasal cavity showing the incisive ducts (ID) and vomeronasal ducts (VND) running independently. The IDs are located ventrally, displaying an elongated and narrow morphology, while the oval-shaped VDs are situated more dorsally. B. Section caudal to image A, where the communication between both ducts is visible (inset), along with their fused course toward the right side of the image. In the central palatal mucosa, the trigeminal nerve and a glandular plexus are observed (arrowhead). C. Higher magnification of the junction between the vomeronasal and IDs as seen in image B. D. View of the two IDs ascending toward the VDs. E. General view of the nasal cavity showing the shift of the IDs from their original ventral position to a more lateral position relative to the VDs, in direct relation to the palatal fissure (arrow). F. General view of the nasal cavity highlighting the morphological context of the incisive and VDs. Vomeronasal cartilage (open arrows in B, C, E). Scale bars: 500 μm (E, F), 100 μm (A, B, D), 50 μm (C).

### Histological study of the accessory and main olfactory bulbs

The accessory olfactory bulb of *Talpa occidentalis* displayed a well-defined structure and spatial organization, as revealed by Nissl staining (Fig. 13). In transverse sections, the AOB was situated dorsomedially with respect to the main olfactory bulb, and in close anatomical proximity to the telencephalon, anterior olfactory nucleus, lateral ventricle, and olfactory nerves (Fig. 13A). A full transverse section through the central part of the AOB revealed its classical laminar architecture, consisting—superficial to deep—of the vomeronasal nerve layer, glomerular layer, mitral-plexiform layer, and granular layer (Fig. 13B). A detailed view of the left AOB highlighted the clear development and separation of the mitral-plexiform and granular layers (Fig. 13C). Further magnification allowed the identification of mitral cells located at the periphery of the glomeruli (Fig. 13D, arrows). In more caudal sections, the AOB exhibited a less conventional organization: numerous glomeruli, periglomerular cells, and mitral-like neurons were arranged in a pattern suggestive of the olfactory limbus, a transitional area of functional and structural interest (Fig. 13E).

**Figure 13.**
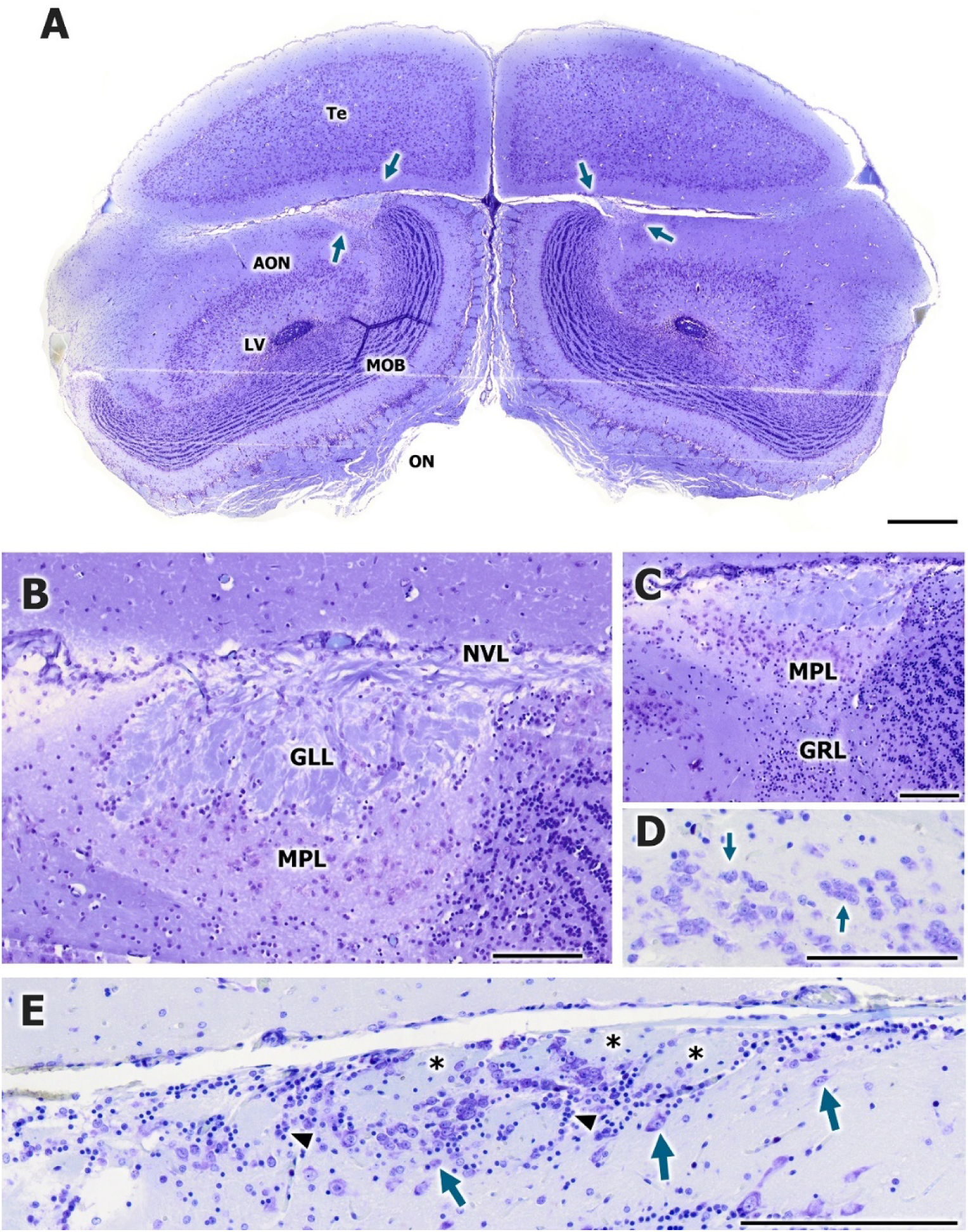
Microscopic study of the accessory olfactory bulb (AOB) of the Iberian mole *Talpa occidentalis* using Nissl staining. A. Transverse section of the olfactory bulb showing the topographic relationships of the AOB (arrows) with the telencephalon (Te) and the main olfactory bulb (MOB). The anterior olfactory nucleus (AON), lateral ventricle (LV), and olfactory nerves (ON) are also visible. B. Complete transverse section of the central portion of the AOB. Its characteristic lamination is distinguishable, consisting from superficial to deep: vomeronasal nerve layer (VNL), glomerular layer (GL), mitral-plexiform layer (MPL), and granular layer (GrL). C. Enlargement of the left accessory olfactory bulb from image A, at a level where the development of the mitral-plexiform and granular layers is clearly visible. D. Higher magnification of mitral cells (arrows) located at the periphery of glomeruli from image E. E. General view of a more caudal transverse section of the accessory olfactory bulb, where glomeruli (*), periglomerular cells, and mitral-like cells (arrows) are observed, forming an atypical organization that may correspond to the olfactory limbus. Scale bars: 100 μm.

The main olfactory bulb (MOB) revealed a well-defined laminar architecture under Nissl staining (Fig. 14). In rostral transverse sections, the MOB displayed a clear and organized stratification across its entire structure (Fig. 14A). In more caudal sections, although the typical layers remained discernible in the ventral region, their development appeared less pronounced compared to the rostral view (Fig. 14B). A detailed examination of the rostral level confirmed the presence of the five canonical layers of the MOB, arranged from dorsolateral to ventromedial: the olfactory nerve layer (ONL), glomerular layer (GL), external plexiform layer (EPL), mitral cell layer (ML), and granular layer (GrL) (Fig. 14C).

**Figure 14.**
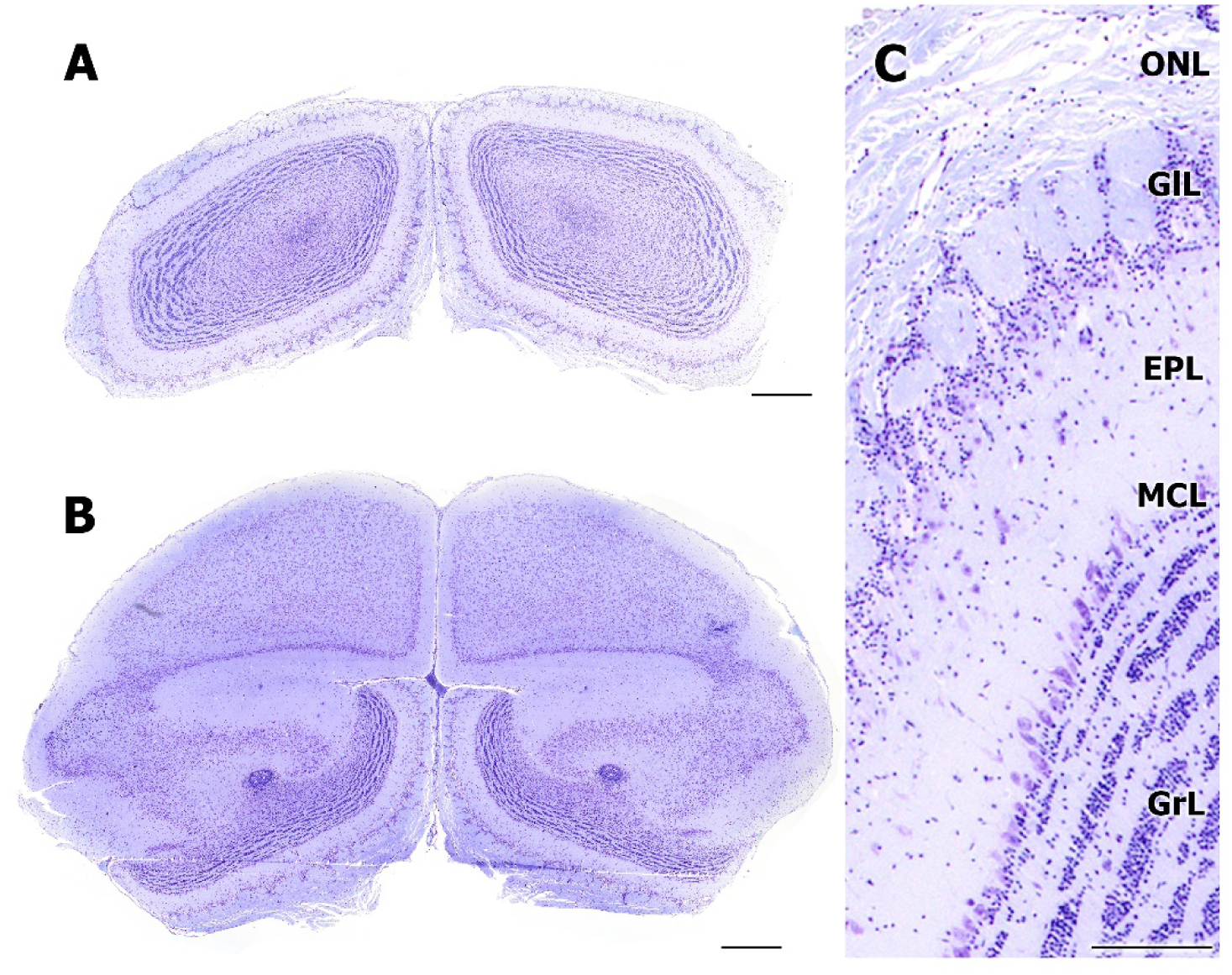
Microscopic analysis of the main olfactory bulb (MOB) using Nissl staining. A. Rostral transverse section of the MOB. The MOB exhibits a prominent laminar organization, clearly visible in this overview. B. Caudal transverse section of the MOB. The characteristic layers of the MOB are visible in the ventral region, though they are less developed than in image A. C. Higher magnification of image A showing the lamination of the MOB, from dorsolateral to ventromedial: olfactory nerve layer (ONL), glomerular layer (GlL), external plexiform layer (EPL), mitral cell layer (MCL), and granular cell layer (GrL). Scale bars: 500 μm (A, B), 125 μm (C).

### Inmunohistochemical study of the accessory olfactory bulb

The accessory olfactory bulb (AOB) of *Talpa occidentalis* showed a diverse pattern of protein expression depending on the immunohistochemical marker used (Fig. 15). Calretinin labeling revealed strong immunoreactivity in the glomerular layer, the vomeronasal nerve layer, and the mitral-plexiform layer, with the most intense staining observed in the granular layer (Fig. 15A). The Gγ8 subunit of G proteins was selectively expressed in the glomeruli, while the nerve, mitral-plexiform, and granular layers remained immunonegative (Fig. 15B). MAP2 expression was concentrated in the mitral-plexiform and granular layers, with weaker staining in the glomerular layer and no immunoreactivity in the vomeronasal nerve layer (Fig. 15C).

**Figure 15.**
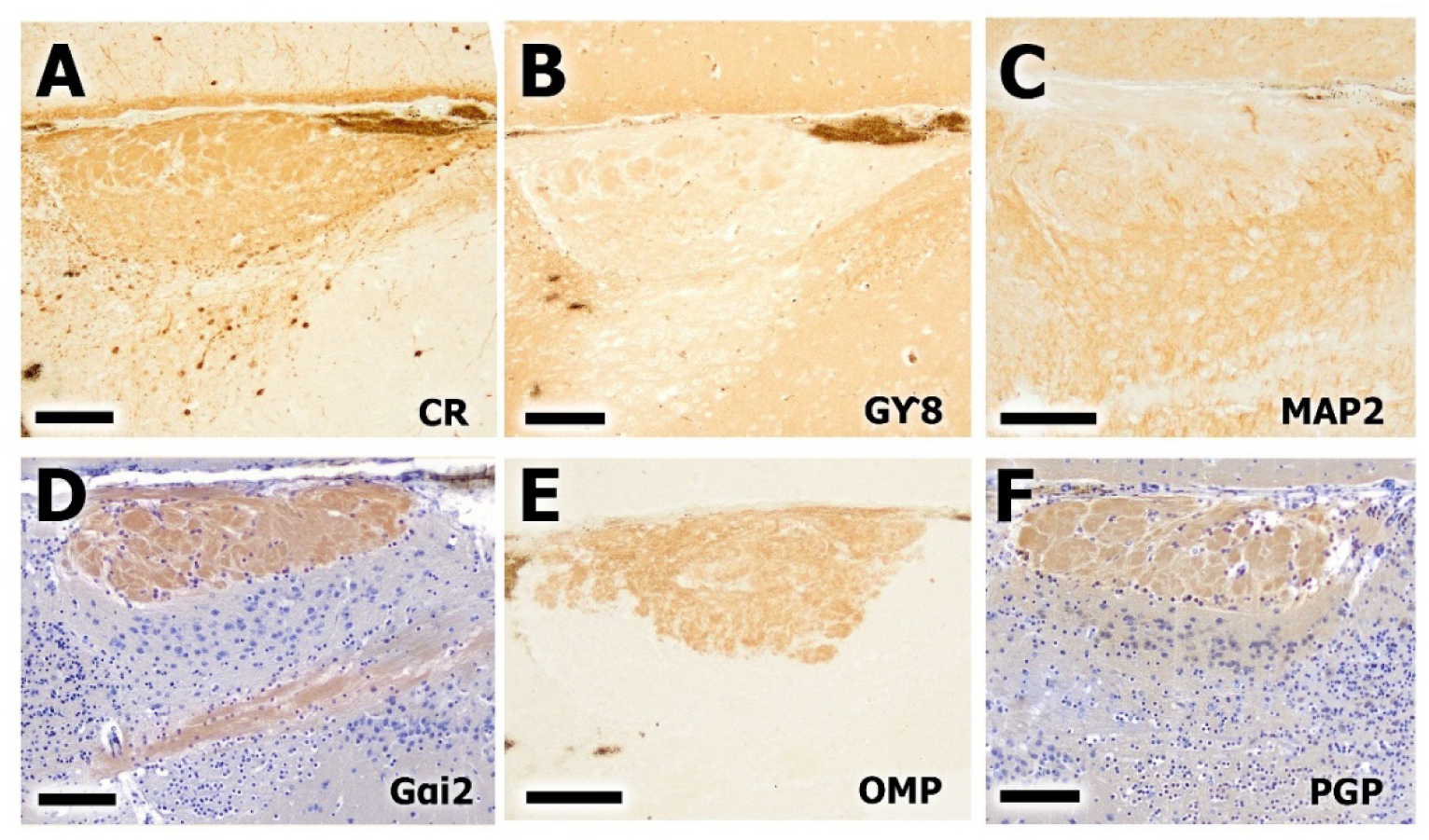
Immunohistochemical analysis of the AOB of the Iberian mole using antibodies against G protein subunits Gαi2 and Gγ8, and the markers CR, MAP2, OMP, and PGP. A. Immunostaining for the calcium-binding protein calretinin. Intense labeling is observed in the glomeruli, the vomeronasal nerve layer, and the mitral-plexiform layer. The most strongly stained cells are located in the granular layer. B. Immunolabeling for the Gγ8 subunit of G proteins. A negative pattern is observed in the nerve, mitral-plexiform, and granular layers, while the glomerular layer shows strong positive labeling. C. Immunostaining for MAP2. This protein is intensely expressed in the mitral-plexiform and granular layers, and weakly in the glomerular layer. The vomeronasal nerve layer is immunonegative. D. Immunohistochemistry for the Gαi2 subunit shows a strongly positive pattern in the nerve and glomerular layers, as well as in the lateral olfactory tract (LOT). Hematoxylin counterstaining. E. Immunostaining for OMP. The anti-OMP antibody reveals strong labeling in the vomeronasal nerve and glomerular layers, with a negative pattern in the remaining layers. F. Immunolabeling for the neuronal marker PGP shows intense positive staining in the nerve, glomerular, and mitral-plexiform layers. Scale bars: 100 μm.

In contrast, the Gαi2 subunit displayed strong labeling in both the vomeronasal nerve and glomerular layers, as well as in the lateral olfactory tract (LOT), suggesting its broader involvement in vomeronasal signaling pathways (Fig. 15D). The OMP marker showed an intense and specific pattern of expression restricted to the vomeronasal nerve and glomerular layers (Fig. 15E). Lastly, the neuronal marker PGP yielded a widespread and robust staining across the nerve, glomerular, and mitral-plexiform layers, highlighting the dense innervation and cellular complexity of the AOB (Fig. 15F).

The immunohistochemical analysis of the MOB revealed distinct and layer-specific patterns of marker expression, reflecting the complex cellular architecture and functional compartmentalization of this structure (Fig. 16). Immunostaining for GAP43, a growth-associated protein linked to axonal remodeling and plasticity, showed selective expression in the granular layer (Fig. 16A). The positive signal was confined to individual neuronal profiles scattered throughout this innermost layer, suggesting active neurite outgrowth or synaptic remodeling among granule cells in the adult MOB. The Gγ8 subunit of G proteins exhibited a markedly compartmentalized pattern of expression. Strong immunoreactivity was observed in the glomerular layer and in the internal plexiform layer, with additional labeling in neuronal populations adjacent to the granular zone (Fig. 16B). In contrast, the nerve layer and mitral cell layer appeared immunonegative. The use of hematoxylin and eosin as a counterstain facilitated the visualization of cytoarchitectural boundaries, underscoring the specificity of the Gγ8 signal within the neuropil-rich regions.

**Figure 16.**
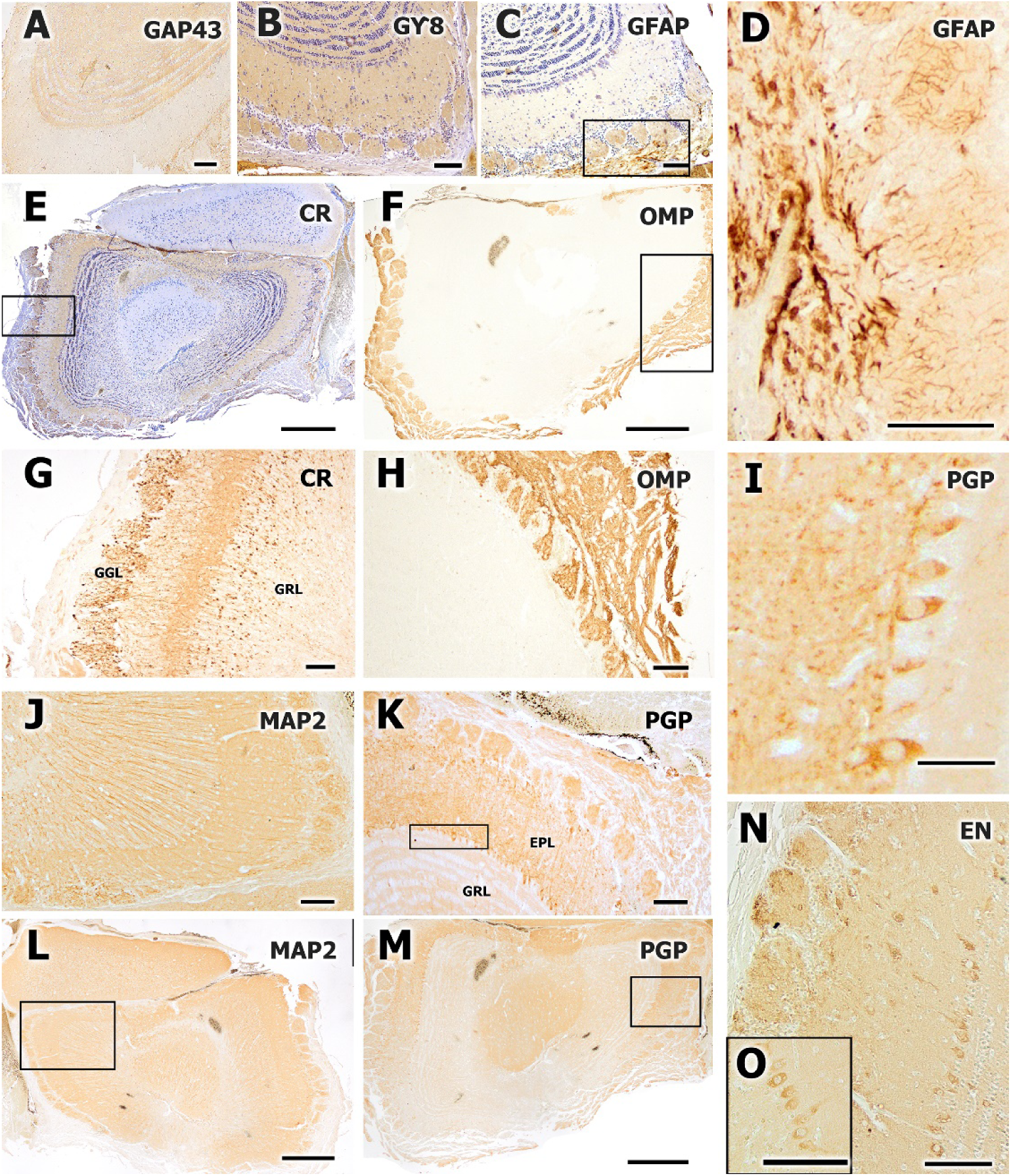
Immunohistochemical analysis of the main olfactory bulb (MOB) using antibodies against GAP43, Gγ8, GFAP, CR, OMP, MAP2, PGP, and EN. A. Immunostaining for GAP43 shows specific labeling of granular cells. B. Immunolabeling for the Gγ8 subunit reveals positive staining in the glomeruli, internal plexiform layer, and in cells bordering the granular bands. Counterstaining with hematoxylin and eosin. C–D. Immunohistochemistry for GFAP. C. General view of the MOB, showing strong labeling in the glomeruli and the nerve layer. D. Higher magnification of the nerve layer and glomeruli from image C, revealing intense labeling of the astrocytes. E, G. Immunostaining for calretinin (CR). E. General view of the AOB and telencephalon, showing variably intense labeling across layers. G. Anti-CR shows strong labeling in glomeruli, mitral cells, and the internal plexiform layer. H. Anti-OMP staining shows intense positivity in the glomerular layer and in its nerve projections throughout the MOB perimeter. J, L. Immunostaining for MAP2. L. General view of the MOB shows positive staining throughout its extent, except for a centrally located unstained groove. J. Enlargement of image L reveals intense staining of axonal projections from the glomeruli to the mitral layer, corresponding to the external plexiform layer. K, M, I. Immunohistochemistry for the neuronal marker PGP. N, O. Anti-Neuron specific enolase strongly immunolabels the mitral cells. Scale bars not specified. Scale bars: 500 μm (E,F, L,M), 100 μm (A,B,C,D,G,H,I,N,O).

Glial fibrillary acidic protein (GFAP) staining provided insights into the distribution of astrocytes across the MOB. A general overview revealed dense labeling in both the glomerular layer and the nerve layer (Fig. 16C). Upon closer examination, intense GFAP immunoreactivity was noted in astrocytic processes encircling the glomeruli and in radial glia within the nerve fiber layer (Fig. 16D), indicating a supportive glial network associated with olfactory sensory input and processing. Calretinin (CR), a calcium-binding protein expressed by specific interneuron populations, displayed variable intensity across MOB layers. In a general view, CR-positive cells were observed throughout the bulb and the adjoining telencephalon (Fig. 16E). At higher magnification, intense immunoreactivity was clearly localized within the glomeruli, the internal plexiform layer, and among mitral cells (Fig. 16G), suggesting that CR may play a role in shaping inhibitory circuits and modulating olfactory signal transmission at multiple levels.

The olfactory marker protein (OMP) showed a highly restricted but robust pattern of labeling. Immunostaining was intense in the glomerular layer and extended along afferent projections that surrounded the entire bulb (Fig. 16H), confirming the widespread and uniform innervation of glomeruli by olfactory receptor neurons. MAP2, a cytoskeletal protein associated with dendritic architecture, exhibited widespread expression throughout the MOB (Fig. 16L). The only exception was a centrally located groove that remained consistently unstained, possibly corresponding to the region of the olfactory peduncle. Higher magnification revealed dense MAP2-positive fibers bridging the glomerular and mitral layers, corresponding to the external plexiform layer (Fig. 16J). This organization reflects the dendritic arborization of mitral and tufted cells.

PGP (Protein Gene Product 9.5), a general neuronal marker, showed broad and intense labeling across multiple layers of the MOB, including the nerve layer, glomeruli, and the mitral-plexiform region (Fig. 16I, K, M). This widespread staining highlights the high density of neuronal elements and the intricate network of afferent and intrinsic fibers present in the bulb. Finally, neuron-specific enolase (NSE) immunostaining provided clear labeling of mitral cells within the MOB (Fig. 16N, O). The signal was robust and localized to large neuronal somata, consistent with the high metabolic demand of these principal projection neurons. The selective NSE expression reinforces the distinct neurochemical identity of mitral cells and their relevance in olfactory signal relay to higher brain centers.

### Lectin histochemical study of the accessory and main olfactory bulb

Histochemical staining of the olfactory bulbs of *Talpa occidentalis* using the lectins PHL, STL, and LEA revealed differential glycosylation profiles between the main and accessory olfactory systems, highlighting both regional and laminar specificity (Fig. 17). PHL histochemistry showed a distinct pattern limited to specific central olfactory projections. In a general brain overview, intense PHL binding was observed in the lateral olfactory tract (LOT), whereas the accessory olfactory bulb showed no detectable labeling (Fig. 17A). This suggests that the glycan targets of PHL are expressed predominantly along the centrifugal olfactory pathways but are absent or underrepresented in the vomeronasal projection areas.

**Figure 17.**
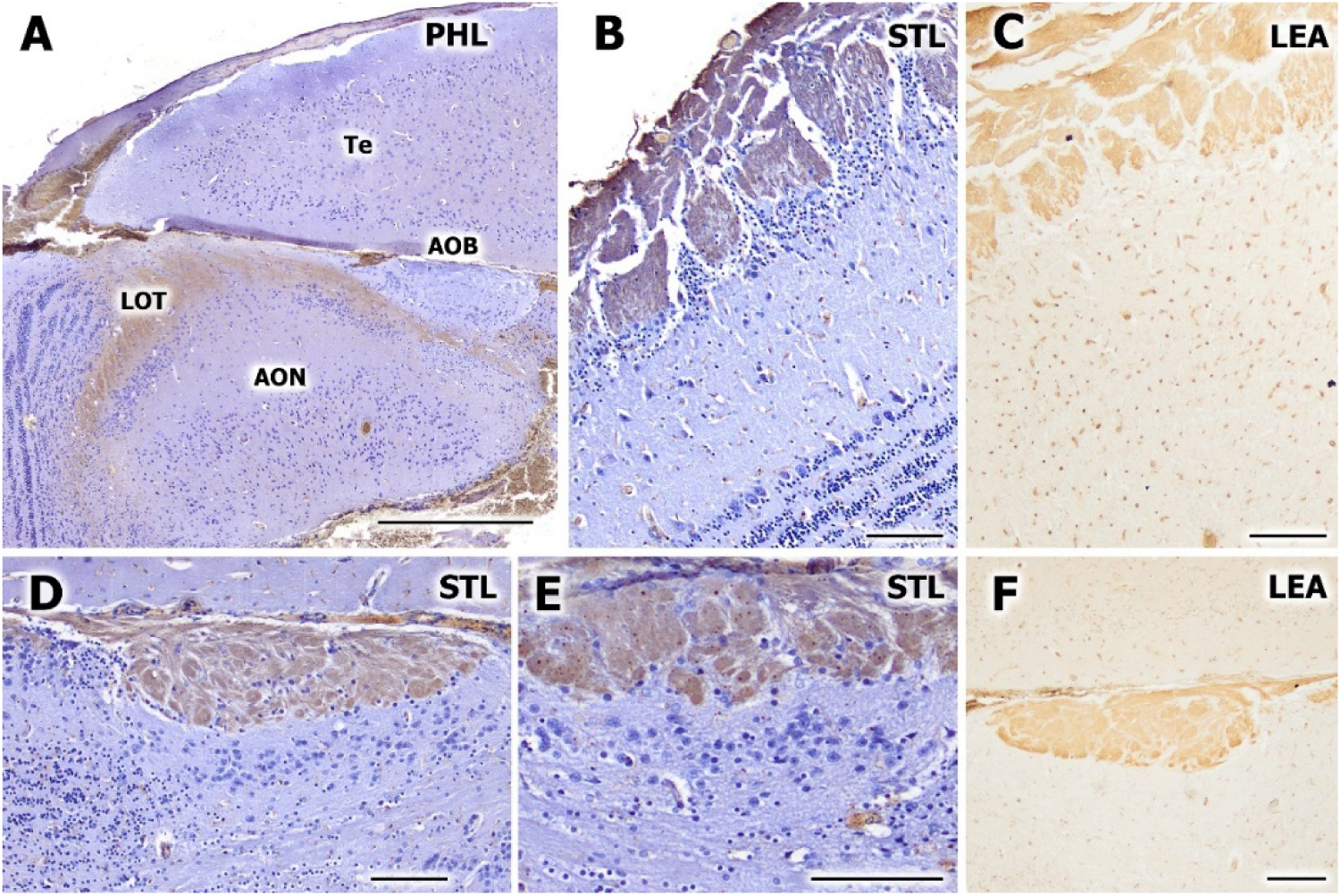
Histochemical analysis of the olfactory bulb of the Iberian mole using the lectins PHL, STL, and LEA. A. General view of the brain showing the accessory olfactory bulb (AOB), the granular layer of the main olfactory bulb (MOB), the telencephalon (Te), and the anterior olfactory nucleus (AON). PHL staining shows intense positivity in the lateral olfactory tract (LOT) and no labeling in the AOB. B, D, and E. STL histochemistry with hematoxylin counterstaining. B. View of the MOB showing very intense staining in the glomeruli and in the tufted cells of the internal plexiform layer. D. View of the AOB with strong positive staining in the glomeruli and in the nerve bundles entering the structure (vomeronasal nerve layer). Hematoxylin counterstaining highlights the different cellular populations forming the laminar architecture of the AOB. E. Higher magnification of the AOB layers. C–F. LEA lectin histochemistry. C. General view of the MOB with intense staining in the glomeruli and nerve layer. F. View of the AOB with very intense staining in the glomerular layer. Scale bars: 500 μm (A), 100 μm (B, C, D, E, F).

STL lectin staining, combined with hematoxylin counterstaining, provided a detailed visualization of both the main and accessory olfactory bulbs. In the MOB, STL exhibited strong labeling within the glomerular layer and in the tufted cells of the internal plexiform layer (Fig. 17B), indicating a high density of STL-binding glycoconjugates in both sensory input zones and principal cell compartments. In the AOB, STL also labeled the glomeruli with high intensity and extended to the vomeronasal nerve layer, where incoming sensory fibers converge (Fig. 17D). The hematoxylin counterstain facilitated the identification of distinct cellular groups, delineating the layered structure of the AOB. A higher magnification of this region allowed clear visualization of its laminar organization, with STL staining marking both glomerular structures and underlying cellular arrangements (Fig. 17E).

LEA staining displayed a broader pattern that encompassed both the main and accessory olfactory structures. In the MOB, LEA bound intensely to the glomeruli and the olfactory nerve layer (Fig. 17C), indicating robust glycan expression in both afferent terminals and periglomerular regions. Similarly, in the AOB, LEA histochemistry produced a very strong signal within the glomerular layer (Fig. 17F), suggesting shared carbohydrate motifs between the main and accessory glomerular circuits despite their distinct functional inputs.

### Double fluorescent labeling study in AOB and MOB with confocal microscopy

Double immunofluorescence staining of the olfactory bulbs of *Talpa occidentalis* revealed distinct and complementary patterns of molecular expression in both the main (MOB) and accessory (AOB) olfactory systems, highlighting the laminar organization and the cellular diversity within these structures (Fig. 18). Immunolabeling with GFAP and GAP43 revealed differential localization of glial and growth-associated elements. In the AOB, GAP43 expression was particularly intense in the glomeruli and in the vomeronasal nerve layer, reflecting high levels of axonal plasticity and sensory input (Fig. 18A). In contrast, GFAP staining was concentrated in the mitral-plexiform and glomerular layers, with additional labeling of interglomerular astrocytes, indicating a supportive glial network. In the MOB, GFAP showed a dense distribution among granular and periglomerular cells, while GAP43 exhibited strong glomerular labeling and moderate signal in the internal plexiform layer (Fig. 18B). A higher magnification of the interglomerular domain confirmed the astrocytic nature of this region, with abundant GFAP-positive processes forming a scaffold between glomeruli (Fig. 18D).

**Figure 18.**
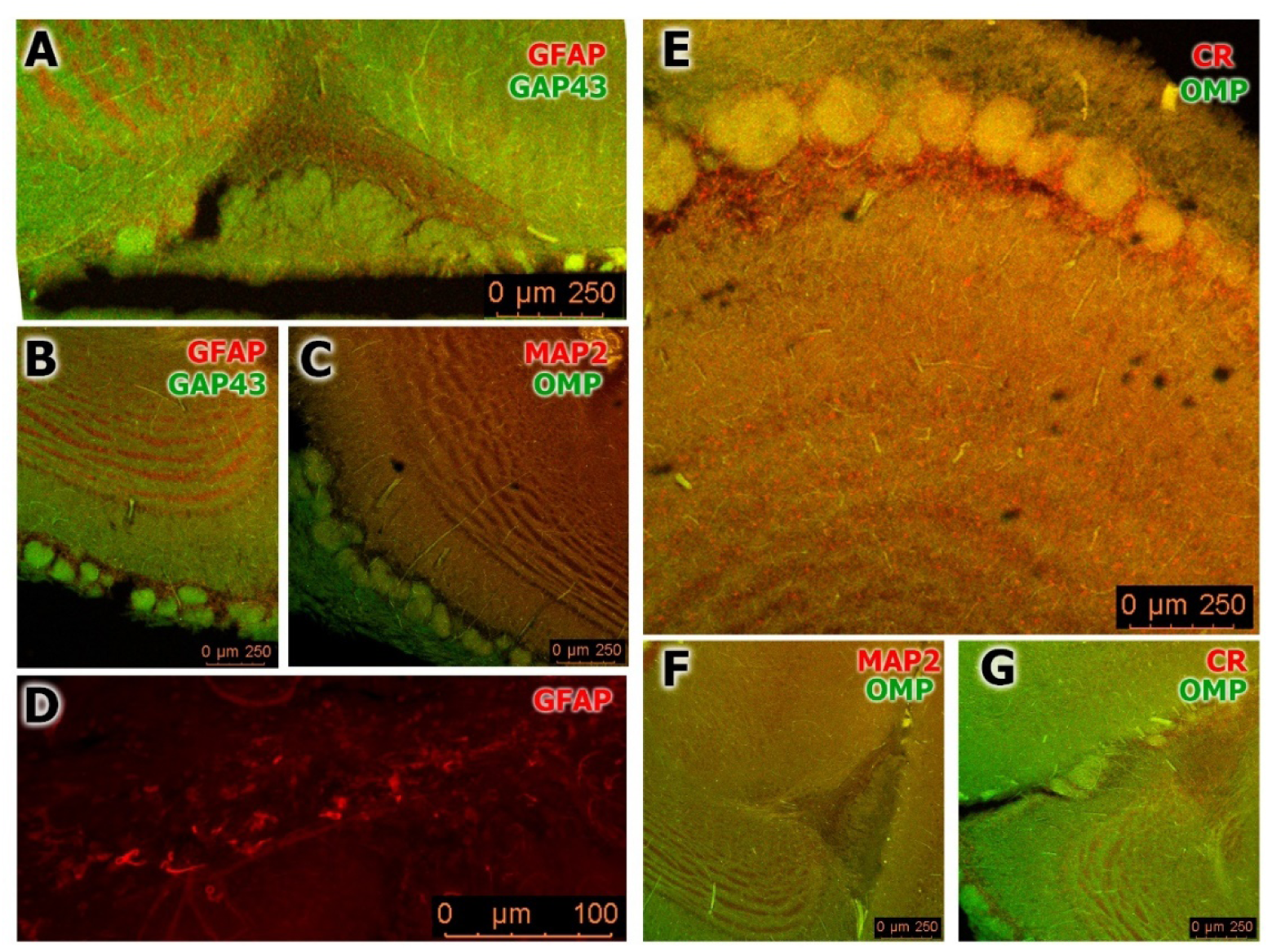
Double immunofluorescence labeling of the olfactory bulbs of the Iberian mole (*Talpa occidentalis*) using antibodies against GFAP, GAP43, OMP, MAP2, and calretinin (CR). A–B. Double immunolabeling for GFAP and GAP43. A. General view of the AOB showing intense GAP43 staining in the glomeruli and vomeronasal nerve layer, while GFAP staining is concentrated in the mitral-plexiform layer and in the glomerular layer, as well as in scattered interglomerular astrocytes. B. General view of the MOB, showing strong GFAP expression in granular cells and periglomerular astrocytes, while GAP43 is intensely expressed in the glomeruli and to a lesser degree in the internal plexiform layer. D. Higher magnification of the interglomerular glial region with abundant GFAP-positive astrocytes. C–F. Double labeling for OMP and MAP2. C. General overview of the MOB showing intense OMP immunoreactivity in the nerve layer, glomeruli, and axonal projections emerging from the glomerular layer. MAP2 shows intense labeling in the granular layer, particularly between clusters of granular cells. Both markers overlap in the mitral and internal plexiform layers, producing a yellow merged signal. F. Transverse view of the AOB showing intense MAP2 staining in the mitral-plexiform layer and predominant OMP expression in the glomerular layer. E–G. Double immunolabeling for OMP and calretinin (CR). E. General view of the MOB showing co-expression of OMP and CR in the glomeruli, mitral-plexiform layer, and granular layer. CR also labels periglomerular astrocytes and some tufted cells in the internal plexiform layer. G. Transverse section of the olfactory bulb identifying the AOB, where strong calretinin staining is observed in the plexiform and granular layers, while OMP shows a more intense signal in the glomeruli. Scale bars: 250 μm (A, B, C, E, F, G), 100 μm (D).

Double labeling for OMP and MAP2 provided insight into the afferent-efferent interactions within the MOB. OMP staining was sharply localized to the olfactory nerve layer, glomeruli, and the axonal projections of sensory neurons (Fig. 18C), whereas MAP2 expression was concentrated in the granular layer, particularly between clusters of granular cells. These two markers overlapped in the mitral and internal plexiform layers, producing a yellow merged signal that marked the integration zone between incoming axons and dendritic arborizations of mitral and tufted cells. In the AOB, MAP2 immunoreactivity was most intense in the mitral-plexiform layer, while OMP staining was predominantly observed in the glomerular layer (Fig. 18F), indicating a similar functional arrangement adapted to vomeronasal input.

Finally, double labeling with OMP and calretinin (CR) further delineated functional subdomains of the olfactory bulbs. In the MOB, both markers were co-expressed in the glomeruli, mitral-plexiform, and granular layers (Fig. 18E), while CR showed particularly strong labeling in periglomerular astrocytes and tufted cells of the internal plexiform layer, suggesting roles in calcium buffering and inhibitory modulation. In the AOB, CR was concentrated in the plexiform and granular layers, whereas OMP maintained its expression in the glomeruli (Fig. 18G), reinforcing the segregation of afferent input and local interneuronal circuitry within the vomeronasal pathway.

### Study of free-floating immunohistochemical labeling in AOB and MOB

The free-floating immunohistochemical analysis of the olfactory bulb of *Talpa occidentalis* using antibodies against calretinin (CR), MAP2, and GFAP provided a detailed visualization of the laminar organization and cellular composition of both the main (MOB) and accessory (AOB) olfactory systems (Fig. 19). Immunolabeling with calretinin revealed an intense and widespread expression across both olfactory bulbs. In a general view of the right hemisphere, strong CR staining was observed in the layers of the MOB and AOB, while adjacent regions of the telencephalon and the anterior olfactory nucleus (AON) remained distinguishable (Fig. 19A). A more detailed view of the AOB showed robust expression of CR throughout most layers, with the exception of the periglomerular zone and the vomeronasal nerve layer, which displayed only faint labeling (Fig. 19C). Further magnification confirmed the presence of intense calretinin expression in periglomerular cells, mitral cells, and the granular layer (Fig. 19D), whereas the internal plexiform layer and glomerular structures showed relatively weaker staining. This pattern supports the presence of calretinin-positive interneurons involved in inhibitory microcircuits, particularly around the glomeruli and principal output layers.

**Figure 19.**
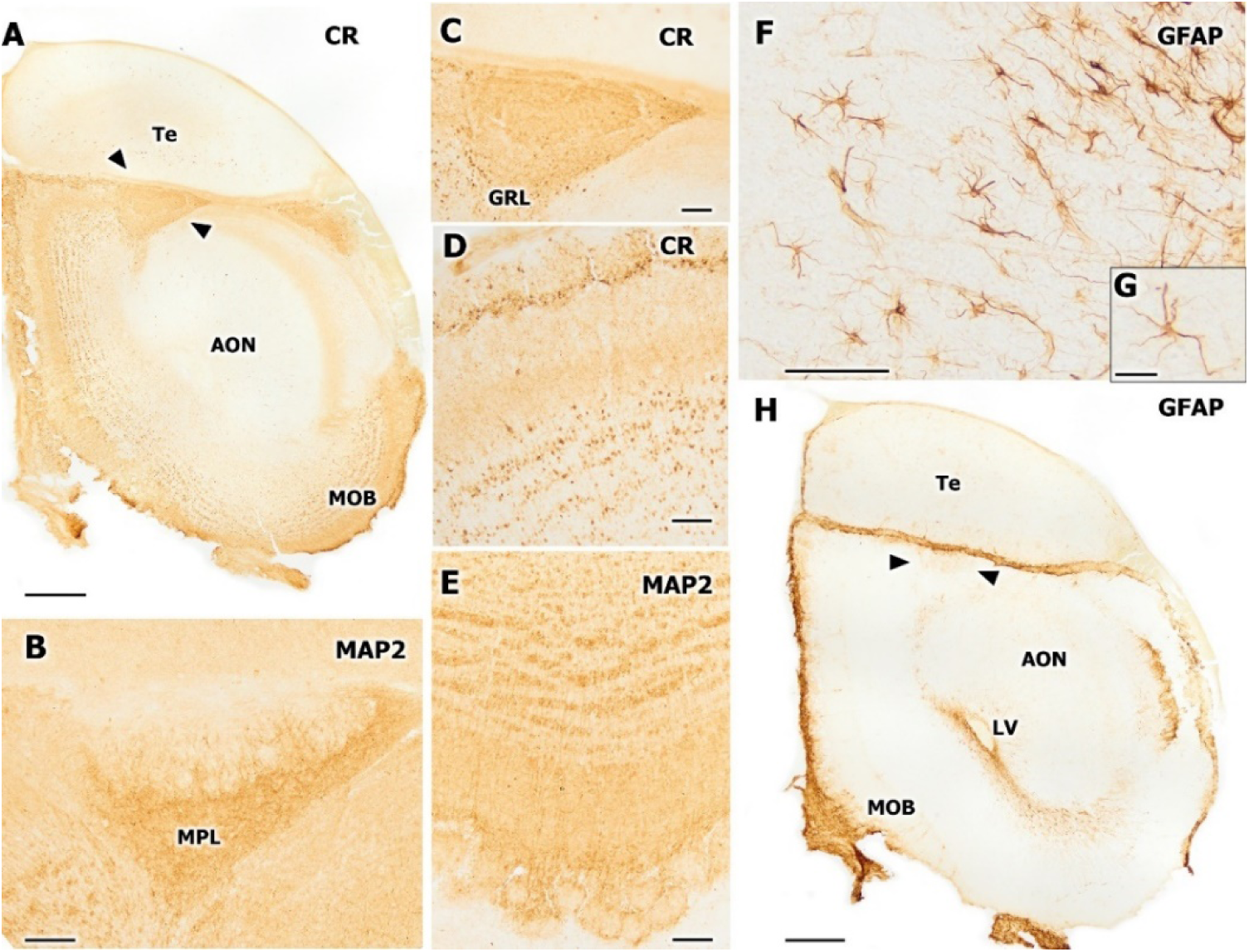
Free-floating immunohistochemical study of the olfactory bulb using antibodies against calretinin (CR), MAP2, and GFAP. A, C, D. Immunolabeling with anti-calretinin antibody. A. General view of the right hemisphere showing intense labeling in the layers of both the MOB and the AOB (arrowheads). The telencephalon (Te) and the anterior olfactory nucleus (AON) are also visible. C. Enlargement of the AOB from image A, showing strong CR expression across all layers, except in the periglomerular region and the nerve layer, where staining is markedly weaker. D. Further enlargement of the AOB highlighting strong CR signal in the periglomerular zone, mitral cells, and granular layer, whereas the internal plexiform layer and glomeruli display weaker staining. B, E. Immunolabeling with anti-MAP2 antibody. B. General view of the AOB showing intense labeling in periglomerular cells, the mitral-plexiform layer (MPL), and the granular layer. In contrast, the nerve layer and glomeruli show minimal or no labeling. E. Laminar structure of the MOB, where MAP2 expression is detected in the granular layer, internal plexiform layer, and in periglomerular cells that outline glomeruli, which themselves show a less intense signal. F, G, H. Immunostaining with anti-GFAP antibody. H. Transverse overview of the OB showing strong labeling in the nerve layer of the MOB and in the lateral ventricle (LV). The telencephalon (Te), anterior olfactory nucleus (AON), and AOB (arrowheads) are also visible. F. Enlargement of the central ventricular zone at a more rostral level than in image H. G. Higher magnification of an astrocyte. Scale bars: 500μm (A,H), 100 μm (B,C,D,E,F), 25μm (G).

The MAP2 marker, which binds to dendritic microtubules, offered complementary information on the distribution of principal and interneuronal dendritic trees. In the AOB, MAP2 was intensely expressed in the periglomerular region, the mitral-plexiform layer, and the granular layer (Fig. 19B), suggesting an active dendritic network supporting sensory integration. The nerve and glomerular layers, by contrast, lacked MAP2 staining, confirming their primarily afferent or non-dendritic composition. In the MOB, MAP2 showed a similar laminar pattern, with strong immunoreactivity in the internal plexiform and granular layers, and in periglomerular cells that outlined each glomerulus (Fig. 19E). The glomerular cores themselves appeared less intensely stained, reinforcing the spatial segregation between afferent synaptic input and postsynaptic dendritic processing zones.

GFAP immunostaining revealed the astroglial scaffold within and around the olfactory bulbs. In a low-magnification transverse view, strong GFAP labeling was visible in the olfactory nerve layer of the MOB and within the walls of the lateral ventricle (Fig. 19H). The image also allowed clear identification of the telencephalon, the anterior olfactory nucleus, and the AOB (arrowheads). In a more anterior section, dense astroglial processes were visualized around the central ventricular zone (Fig. 19F), suggesting radial glial arrangements or ependymal-associated glia. A close-up view of an individual astrocyte (Fig. 19G) revealed a typical multipolar morphology with extended processes, confirming the robust astrocytic presence in periventricular and periglomerular domains.

## Discussion

Sociosexual behaviors, which are essential for the establishment of deep inter-individual relationships within a species, as well as for interspecies survival-related instincts, are primarily mediated through the vomeronasal system (VNS). As such, understanding its structure and function is critical for elucidating these interactions and applying this knowledge in diverse fields, including animal husbandry, endangered species conservation, and agricultural pest control (Dehnhard, 2011). To date, the most extensive body of literature on the VNS—spanning morphology, neurochemistry, physiology, genomics, and experimental models—has focused on rodents (rats and mice), due to their historical and practical value in laboratory settings (Murata et al., 2024b). In recent years, significant advances have also been made in lagomorphs, which now serve as prominent models owing to their well-developed vomeronasal system (Villamayor et al., 2024). Despite this, the VNS is one of the most evolutionarily diverse sensory systems. It has undergone substantial modifications across species and exhibits markedly different evolutionary trajectories even among closely related taxa. This evolutionary plasticity underscores the difficulty of extrapolating findings from one species to another and reinforces the necessity of species-specific investigations (Silva and Antunes, 2017).

Our study addresses this gap by examining the VNS in *Talpa occidentalis*, a species within the order Eulipotyphla (formerly Insectivora), for which prior data are scarce. While previous studies have focused on Erinaceidae and Soricidae families, none has investigated the family Talpidae. The Iberian mole, *T. occidentalis*, is endemic to the Iberian Peninsula and presents a unique opportunity for VNS analysis due to its fossorial lifestyle and sensory adaptations. The Iberian mole (*Talpa occidentalis*) exhibits a series of anatomical and ecological traits that render the study of its vomeronasal system (VNS) particularly distinctive. As a strictly subterranean species, its sensory systems have undergone marked adaptations to its underground environment. Visual perception is extremely limited, restricted to the detection of light stimuli (Carmona et al., 2010), and auditory capabilities are similarly reduced, being responsive only to strong vibrations or loud sounds. These sensory constraints suggest a compensatory enhancement of alternative sensory modalities, notably the vomeronasal, olfactory, and tactile systems.

As previously outlined, the vomeronasal system comprises three main components: the vomeronasal organ (VNO), the vomeronasal nerves, and the accessory olfactory bulb (AOB) (Vaccarezza et al., 1981). Histological examination of decalcified nasal cavity sections from *T. occidentalis* enabled the clear identification of paired vomeronasal organs situated bilaterally along the vomer bone. These structures exhibited a distinct anatomical organization compared to that described in other small mammals. In most small mammalian species, the VNO is encased by a J-shaped bony or cartilaginous capsule that serves to protect the vomeronasal duct (Kondoh et al., 2017b; Ortiz-Leal et al., 2020; Torres et al., 2020). However, in *T. occidentalis*, although the vomeronasal capsule was composed of hyaline cartilage— similar to that described in the African pygmy hedgehog (*Atelerix albiventris*) (Kondoh et al., 2021)—its configuration did not conform to the classical morphology. Instead, the cartilage was exclusively located in the space between the maxillary and vomer bones, providing ventral support to the VNO. Moreover, unlike in other species, the vomeronasal cartilage in *T. occidentalis* was only present in a limited number of central sections, rather than forming a continuous capsule along the full length of the duct and its associated parenchyma.

Histological analysis of the vomeronasal duct using hematoxylin-eosin staining revealed a circular ductal morphology, similar to that reported in the African pygmy hedgehog (*Atelerix albiventris*), (Kondoh et al., 2021) and notably different from the crescent-shaped lumen typically observed in reference species such as rodents (Salazar and Sánchez Quinteiro, 2003; Mechin et al., 2021) and lagomorphs (Elgayar et al., 2014). Upon higher magnification, the duct in *T. occidentalis* was found to be lined by a single, continuous, and uniform epithelium extending along its entire perimeter. To our knowledge, this homogeneous epithelial arrangement is unique among vertebrates. In the most extensively studied species (rodents and lagomorphs), the vomeronasal duct is consistently lined by two distinct epithelial types: a sensory epithelium and a non-sensory (or respiratory-like) epithelium, distributed in varying proportions (Garrosa and Coca, 1991; Mendoza, 1993). This dual epithelial organization has been documented in all species examined to date, including the hedgehog. In *A. albiventris*, for instance, the rostral region of the vomeronasal organ is mainly lined by sensory epithelium, the caudal region by non-sensory epithelium, and both epithelial types coexist in the central portion of the duct (Kondoh et al., 2021).

In *T. occidentalis*, the vomeronasal parenchyma appears poorly developed, as indicated by the near absence of prominent blood vessels or arteries, which were notably small in diameter. Vomeronasal nerves were restricted to the dorsolateral region of the parenchyma adjacent to the nasal septum, and a dense glandular plexus was observed in the same area. The most striking and potentially limiting feature, uptake of semiochemicals. This mechanism, known as the vomeronasal pump, relies on one or more large-caliber vessels surrounding the lumen of the vomeronasal duct, which generate a negative pressure gradient to draw chemical stimuli into the organ (Meredith et al., 1980; Meredith, 1994). In rodents such as rats and mice, a large vein running parallel to the duct supports the function of this pump (Cantó Soler and Suburo, 1998; Ignacio Salazar et al., 1998; Dennis et al., 2020; Hamacher et al., 2024; Ruiz-Rubio et al., 2024b). In *T. occidentalis*, however, no comparable vascular structure was identified, suggesting that such a mechanism is either absent or functionally irrelevant. The most plausible explanation for this deficiency lies in two anatomical features: (1) the unusually close topographic relationship between the vomeronasal duct and the nasal cavity, which could allow for direct passive entry of semiochemicals, and (2) the presence of a sensory neuroepithelium that lines the entire duct from its rostral extent, thus maximizing contact surface for chemical detection. The absence of a vomeronasal pump may also explain the loss of the classical J-shaped morphology of the cartilaginous capsule—a highly unusual trait. In species where negative pressure is generated, a rigid capsule may serve to prevent parenchymal collapse. In *T. occidentalis*, by contrast, the absence of this mechanical stress would negate the need for such structural reinforcement. To our knowledge, no other species has been reported to lack a vomeronasal pump, making the Iberian mole a uniquely divergent model for studying vomeronasal function.

To assess the functional capacity of the vomeronasal system (VNS), the morphological analysis was complemented with an immunohistochemical study using an extensive panel of antibodies. Among the most informative markers were anti-Gα0, anti-Gαi2, anti-Gγ8, anti-PGP, anti-OMP, anti-GAP43, and anti-calretinin (CR). Particular attention was given to the expression patterns of Gα0 and Gαi2 subunits, as they are directly associated with the two main vomeronasal receptor families: V2R and V1R, respectively—V2Rs coupling to Gα0 (Matsunami and Buck, 1997), and V1Rs to Gαi2 (Dulac and Axel, 1995). The presence or absence of these receptor families has served as the basis for classifying mammals into two vomeronasal system models. The segregated model, typical of Afrotheria, features the co-expression of both receptor families—V1R and V2R—distributed in distinct zones of the vomeronasal epithelium and the accessory olfactory bulb (Jia and Halpern, 1996). In contrast, the uniform model, commonly observed in Laurasiatherian species, exhibits functional expression of only V1R receptors (Takigami et al., 2000). Given that the family Talpidae, to which *T. occidentalis* belongs, occupies a phylogenetic position between these two mammalian superorders, the study of G-protein subunit distribution in this species is of particular relevance.

In *T. occidentalis*, Gα0 immunoreactivity was primarily localized to the apical region of the sensory epithelium and to selected nerve bundles. In contrast, Gαi2 and Gγ8 exhibited overlapping labeling in the apical knobs of bipolar sensory neurons, as well as in their ciliary projections and axonal extensions toward the brain. This pattern, though divergent from that reported in rodent models—where Gα0 and Gαi2 display complementary expression in the somata of vomeronasal sensory neurons (Salazar and Sánchez Quinteiro, 2003)—is consistent with the hypothesis that vomeronasal receptors interact with semiochemicals primarily at the apical knobs and ciliary domains (Tomiyasu et al., 2017). The relatively higher intensity of Gαi2 and Gγ8 labeling, compared to that of Gα0, suggests a predominance of Gαi2-mediated signaling in *T. occidentalis*, indicative of an enrichment in V1R-type receptors. On the basis of these expression profiles, the vomeronasal system of *T. occidentalis* would be best classified within the segregated model of mammalian VNS organization.

Regarding the remaining antibodies, a notable overlap was observed in the labeling patterns of OMP and calretinin (CR) within the neuronal somata of bipolar vomeronasal sensory neurons, particularly in the dorsolateral region of the duct. These patterns are atypical when compared to other species, where CR immunoreactivity is usually confined to the apical knobs of sensory neurons (Torres et al., 2023a), and OMP staining is generally more intense, reflecting a higher degree of neuronal maturation (Albeanu et al., 2018). In contrast, GAP43 immunoreactivity was restricted to the apical knobs and their ventral projections, showing intense labeling limited to these domains. Interestingly, PGP exhibited a robust labeling pattern that combined features of OMP, CR, and GAP43, with strong immunopositivity in the somata of bipolar neurons—especially in the dorsolateral region—and in their apical knobs. These findings support the interpretation that the entire perimeter of the vomeronasal duct in *T. occidentalis* is lined with sensory, non-respiratory epithelium. Moreover, PGP also labeled the axonal projections of bipolar neurons, reinforcing the functional integrity of this receptor epithelium.

To further characterize the receptor environment, lectin histochemistry was performed. Lectins LEA, SBA, ECL, and VVA exhibited strong labeling in the sensory cilia, indicating a high concentration of glycoconjugates in these receptor structures and suggesting a potential role for these biomolecules in vomeronasal chemosignal transduction. Lectins DBA, STL, and LCA revealed specific binding patterns within the sensory epithelium, with SBA particularly highlighting the somata of bipolar neurons. The lectin PHL displayed a highly specific pattern, labeling only the vomeronasal parenchymal glands and the basal cells of the sensory epithelium.

Following the comprehensive analysis of the vomeronasal organ, an equivalent study was conducted on the accessory olfactory bulb (AOB), the central processing center for vomeronasal input. In *T. occidentalis*, the AOB displayed the characteristic lamination seen in well-developed vomeronasal systems (Larriva-Sahd, 2008; Suárez et al., 2011b), including a prominent glomerular layer distinctly bordered by periglomerular cells, an extensive mitral-plexiform layer, and a densely cellular granular layer forming an inverted pyramidal configuration.

A remarkable feature of the AOB in *T. occidentalis* is its relatively small size in proportion to the brain, combined with a high degree of laminar differentiation. This contrasts with the general rule observed in mammals, whereby poorly laminated AOBs (e.g., in ferrets: (Kelliher et al., 2001); and dogs: (Nakajima et al., 1998; Salazar et al., 2013)) are typically associated with reduced size, while larger AOBs exhibit clearer lamination. *T. occidentalis* thus represents a rare exception to this pattern, comparable only to species such as the meerkat (*Suricata suricatta*) (Torres et al., 2021), although in that case the lamination is not as well-defined as in the Iberian mole.

To delineate and characterize the different AOB layers, a battery of antibodies was employed, including anti-Gα0, anti-Gαi2, anti-Gγ8, anti-MAP2, anti-CR, anti-OMP, and anti-PGP. Gα0 and Gγ8 displayed similar patterns, being restricted to selected glomeruli and at low intensity, whereas Gαi2 showed strong immunoreactivity in the glomerular layer and the lateral olfactory tract (LOT). These results support a preliminary classification of *T. occidentalis* within the segregated vomeronasal model. However, the relatively low expression of Gα0 in the glomerular layer suggests that further immunohistochemical verification on additional AOB and VNO sections is warranted. Additionally, PGP, OMP, and CR exhibited overlapping immunoreactivity in the glomerular layer, while CR and PGP also co-labeled neurons in the mitral-plexiform layer. Among these, CR was particularly notable for its strong labeling of granular cells, a pattern consistent with findings in *Erinaceus europaeus* (Briñón et al., 2001). The dendritic marker MAP2 highlighted a dense dendritic plexus in the mitral-plexiform layer and clearly defined the glomerular border.

Lectin histochemistry of the AOB yielded more limited results: despite the use of a broad lectin panel, only STL and LEA produced specific labeling, both localized to the glomerular layer. This is unexpected, given that most mammalian species exhibit more consistent glycosylation patterns in the AOB despite interspecific structural diversity (Ichikawa et al., 1994; Shapiro et al., 1995; Salazar et al., 2001; Shin et al., 2017; Tomiyasu et al., 2018). The atypical lectin-binding pattern observed in *T. occidentalis* suggests a unique glycoconjugate profile that merits further investigation. The co-localization of STL and LEA in glomerular territories also points to a potential role for these glycoconjugates in synaptic interactions between first- and second-order neurons in the vomeronasal pathway.

In conclusion, the vomeronasal system of *Talpa occidentalis* displays a unique combination of anatomical, histological, and neurochemical features that distinguish it from all previously described mammalian models. The absence of a functional vomeronasal pump, the presence of a continuous sensory epithelium lining the entire duct, the loss of the classical J-shaped cartilage, and the distinctive immunohistochemical and lectin binding patterns all point to an alternative functional strategy for chemosensory processing in this fossorial species. These adaptations likely reflect evolutionary pressures associated with its subterranean lifestyle and reduced reliance on visual and auditory cues. The findings not only expand our understanding of vomeronasal diversity within Eulipotyphla but also challenge assumptions derived from rodent-based models, reinforcing the need for broader comparative analyses. *T. occidentalis* thus emerges as a valuable model for exploring the evolutionary plasticity and ecological specialization of the vomeronasal system in mammals.

## AUTHOR CONTRIBUTIONS

Conceptualization, G.G.H., M.G.A.E., P.S.Q., and I.O.L.; Methodology, G.G.H., A.M.A., M.G.A.E., P.S.Q., and I.O.L.; Investigation, G.G.H., A.M.A., M.G.A.E., A.V.C., P.S.Q. and I.O.L.; Resources, A.V.C., J.L.R., and P.S.Q.; Writing – Original Draft Preparation, G.G.H.; Writing – Review & Editing, G.G.H., P.S.Q, and I.O.L.; Supervision, P.S.Q. and I.O.L.; Project Administration, P.S.Q. and I.O.L.; Funding Acquisition, P.S.Q.

## INSTITUTIONAL REVIEW BOARD STATEMENT

Not applicable, as all the animals employed in this study died by natural causes.

## INFORMED CONSENT STATEMENT

Not applicable, as this research did not involve any humans.

## DATA AVAILABILITY STATEMENT

All relevant data are within the manuscript and are fully available without restriction.

